# Chromatin remodeler Brahma safeguards canalization in cardiac mesoderm differentiation

**DOI:** 10.1101/2020.06.03.132654

**Authors:** Swetansu K. Hota, Andrew P. Blair, Kavitha S. Rao, Kevin So, Aaron M. Blotnick, Ravi V. Desai, Leor S. Weinberger, Irfan S. Kathiriya, Benoit G. Bruneau

## Abstract

Differentiation proceeds along a continuum of increasingly fate-restricted intermediates, referred to as canalization^1–4^. Canalization is essential for stabilizing cell fate, but the mechanisms underlying robust canalization are unclear. Here we show that deletion of the BRG1/BRM-associated factor (BAF) chromatin remodeling complex ATPase gene *Brm* (encoding Brahma) results in a radical identity switch during directed cardiogenesis of mouse embryonic stem cells (ESCs). Despite establishment of well-differentiated precardiac mesoderm, *Brm*-null cells subsequently shifted identities, predominantly becoming neural precursors, violating germ layer assignment. Trajectory inference showed sudden acquisition of non-mesodermal identity in *Brm*-null cells, consistent with a new transition state inducing a fate switch referred to as a saddle-node bifurcation^3,4^. Mechanistically, loss of *Brm* prevented de novo accessibility of cardiac enhancers while increasing expression of the neurogenic factor POU3F1 and preventing expression of the neural suppressor REST. *Brm* mutant identity switch was overcome by increasing BMP4 levels during mesoderm induction, repressing *Pou3f1* and re-establishing a cardiogenic chromatin landscape. Our results reveal BRM as a compensable safeguard for fidelity of mesoderm chromatin states, and support a model in which developmental canalization is not a rigid irreversible path, but a highly plastic trajectory that must be safeguarded, with implications in development and disease.

## Main Text

Our previous studies indicated the prevalence of BRM in the cardiomyocyte-enriched chromatin remodeling complex BAF170^5^. BRM has been reported to be dispensable for mouse development^6^, but it is implicated in human developmental syndromes^7,8^, mouse skeletal muscle function^9^, and several cancers^10,11^, and can partly compensate for loss of BRG1^12–16^. To determine the role of BRM in cardiac differentiation, we deleted *Brm* in ESCs and directed cardiac differentiation (Fig. 1a, Extended data Fig. 1a). *Brm*^*−/−*^ cells failed to generate beating cTnT+ cardiomyocytes (Extended Data movies. 1-3) and as measured by immunofluorescence (Fig. 1b) and flow cytometry (Fig. 1c). This was confirmed in three independent *Brm*^*−/−*^ lines, while two heterozygous lines from the same set of clones differentiated well. RNA-seq during the differentiation time course showed that at D4, mesoderm gene expression was unaffected in *Brm*^*−/−*^ cells, while D6 (cardiac precursor, CP) and D10 (cardiomyocyte, CM) gene expression was significantly altered (FDR<0.05, ±2-fold change) (Extended data Fig. 1b, c). At D6, several important cardiac TFs were not induced in *Brm*^*−/−*^ cells (*Isl1*, *Hand2*, Nkx2-5, *Mef2c*, *Tbx20*), whereas osteoblast- and neural-associated TFs were upregulated (*Tcf15, Sox2*). At D10, when WT cells had reached the beating cardiomyocyte stage, *Brm*^*−/−*^ cells completely failed to activate cardiac genes and instead expressed genes associated with neural (*Ascl1*, *Pax6*, *Neurod1*, *Neurog1*, *Olig2* and *Sox2*) or other (e.g. erythrocyte *Gata1*, *Tal1*) cell types (Fig. 1d).

**Fig. 1.**
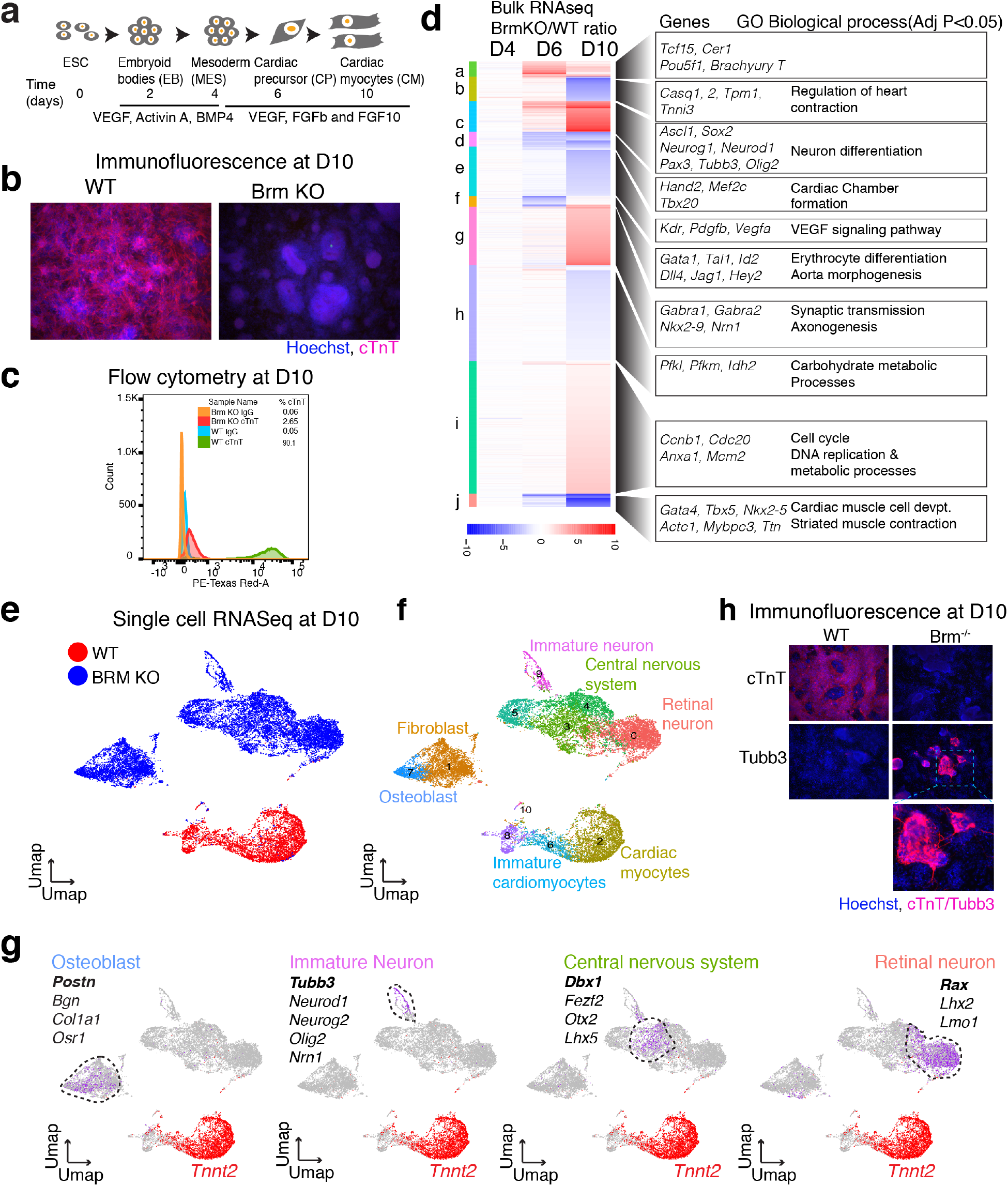
BRM activates cardiac gene expression programs and represses neural genes during directed cardiomyocyte differentiation. Cardiac differentiation scheme (**a**), estimation of cardiac myocyte at D10 of differentiation by immunofluorescence (**b**) and flow cytometry (**c**) of cardiac Troponin T. **d**, Bulk RNAseq analysis of WT and BRM KO cells at mesoderm (D4), cardiac precursor (D6) and cardiomyocyte (D10) stages of differentiation. Counts per million (CPM) average of three replicates were plotted as a ratio of WT over KO. Gene Ontology (GO) biological process enrichment were determined by GOElite. Single cell RNAseq gene expression projected on a UMAP space for WT and BRM KO at D10 (**e**), marker genes highlighted (**f**) and cell types inferred (**g**). **h**, immunofluorescence of cardiac Troponin T and pan-neural progenitor marker TUBB3 (TUJ1) at D10 of cardiac differentiation.

### BRM safeguards CP differentiation to CM and represses neural and other gene programs

The drastic gene expression changes in *Brm*^*−/−*^ cells upon cardiac differentiation suggested either 1) the formation of new non-cardiac populations or 2) that a relatively homogeneous population activated normally mutually-exclusive expression modules. To differentiate between these scenarios, we performed single cell RNA-seq on 21,991 WT and *Brm*^*−/−*^ cells at D10 of differentiation using the 10X Genomics dropseq platform. Uniform Manifold Approximation and Projections (UMAP)^17^ plots showed that *Brm*^*−/−*^ cells at D10 were radically distinct from their WT counterparts (Fig. 1e), lacked expression of cardiac genes, and clustered in multiple sub-populations. These included cells with signatures of neural stem cells (*Sox2, Sox9, Neurod1*, *and Ascl1*), neural progenitors or immature neurons (*Dcx, Otx2, Gap43, Tubb2b, Tubb3*), glial (*Gfap, Olig2*) and Schwan cells (*Gap43*), and retinal neuronal precursors (*Rax, Lhx2, Lmo1*) (Figs. 1f, g, Extended Data Fig. 1d). We also identified a cluster of cells expressing markers of osteoblast development (*Postn, Bgn, Col1a1, Fbln2*, *Twist2*), indicating that some *Brm*^*−/−*^ cells adopt non-cardiac mesodermal fates. Immunofluorescence showed TUBB3+ staining in D10 *Brm*^*−/−*^ cells, displaying neuron like outgrowths (Fig. 1h). Notably, no other mesodermal or ectodermal derivatives, nor endodermal cell types, were observed, indicating that BRM deletion induced a specific fate switch. Loss of BRM did not seem to affect directed neuronal precursor differentiation (Extended Data Fig. 1e).

### BRM controls the mesoderm to cardiac precursor transition

To elucidate the events underlying the anomalous differentiation in absence of BRM, we examined the time courses of differentiation of WT and *Brm*^*−/−*^ cells by single cell RNA-seq (Fig. 2a). Consistent with bulk RNA-seq, D4 (mesoderm) *Brm*^*−/−*^ cells were statistically similar to WT cells and both occupied the same UMAP space (Fig. 2b-d). *Brm*^*−/−*^ cells segregated slightly based on modest (less than 2-fold) changes in expression of a few genes (*Mesp1, Lhx1, Fn1, Rps28*) (Supplementary Sheet 1). In sharp contrast, at D6 *Brm*^*−/−*^ clustered separately from WT cells (Fig. 2b). Most D6 WT cells expressed well-defined CP markers (*Smarcd3, Mef2c, Hand2*; Fig. 2d, Extended Data Fig. 2a), whereas *Brm*^*−/−*^ cells mostly expressed genes involved in neural lineages (*Gbx2, Sox2, Irx3, Crabp1, Crabp2* and *Prtg;* Fig. 2d, Extended Data Fig. 2a). The few D6 WT and *Brm*^*−/−*^ cells that clustered together expressed markers of hematopoietic lineages (*Gata1, Klf1*, *Hbb1*; Supplementary Sheet 2), suggesting a low level of BRM-independent hematopoietic differentiation. As expected, D10 WT and *Brm*^*−/−*^ cells clustered largely in different UMAP space (Fig. 2b-d). Our time course therefore indicates a crucial early role of BRM immediately following cardiac mesoderm formation. Partition Based Graph Abstraction^18^ revealed genotype-dependent connectivity (Fig. 2e-g). WT and *Brm*^*−/−*^ cells at D4 predominantly connected with their respective D6 and D10 genotype-specific clusters. A small percentage of WT and *Brm*^*−/−*^ D6 cells connected to clusters 5 and 12 forming hematopoietic and endothelial clusters (Fig. 2e-g). To further assess the differentiation paths, we built differentiation trajectories in the form of a branching tree using URD^19^. *Pou5f1*+ −WT and *Brm*^*−/−*^ clusters were selected as the root, while D10 clusters were defined as tips. Only WT cells progressed in stepwise pseudotime to CPs and their derivatives. In contrast, *Brm*^*−/−*^ cells directly transitioned from mesoderm to non-cardiac neural lineages after D4 (Fig. 2h, i, Extended Data Fig. 2b, c).

**Fig. 2.**
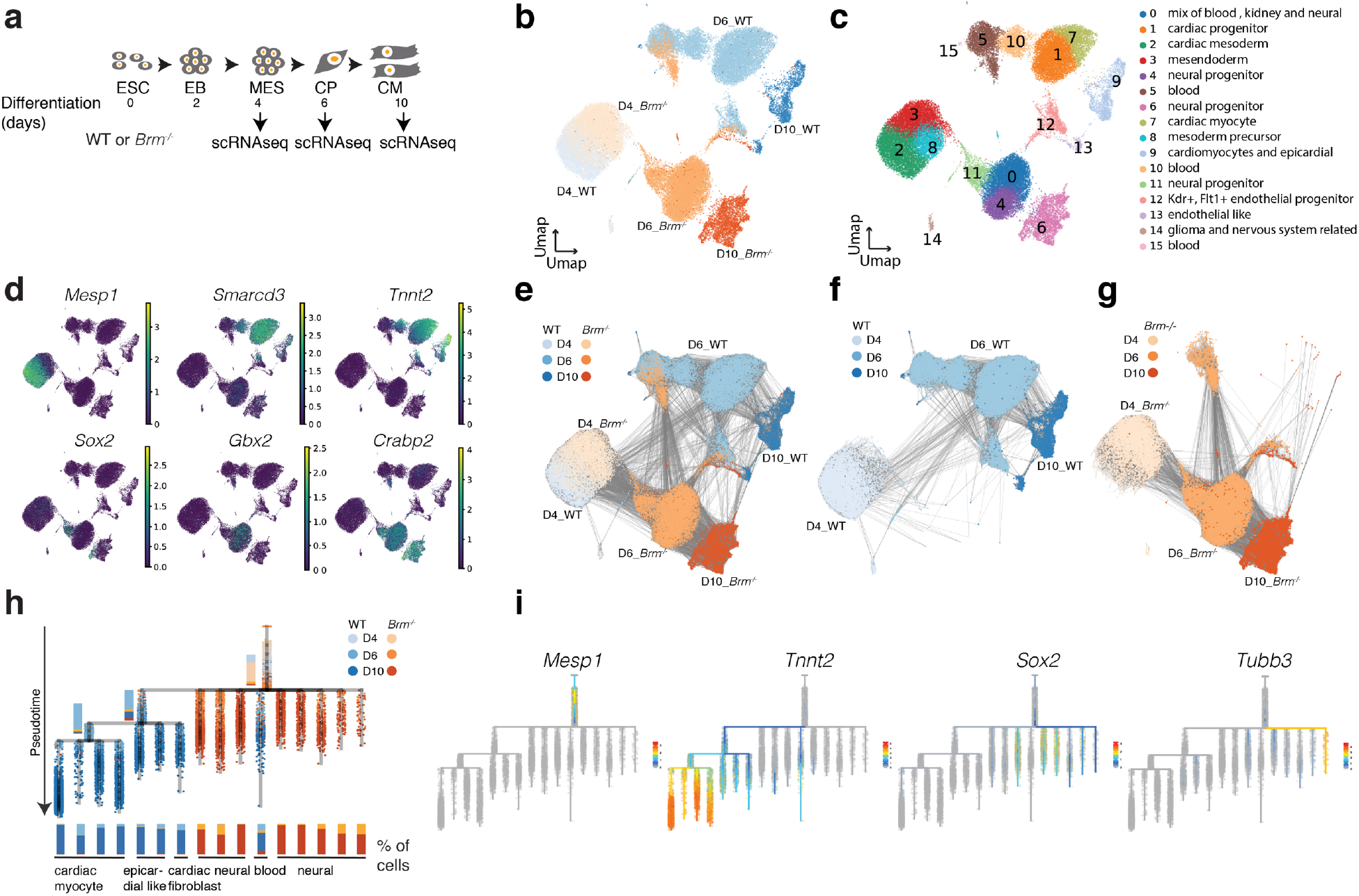
Loss of BRM leads to acquisition of neural fate after pre-cardiac mesoderm formation. **a**, Scheme of cardiac differentiation and time of scRNAseq. **b-d**, Processed scRNAseq data projected on a UMAP space showing genotypes and days of differentiation (**b**), clusters with inferred cell types (**c**), and gene expression feature plots displaying expression of cardiac and neural genes along differentiation (**d**). **e-g**, Partition-based graph abstraction (PAGA) showing connectivity of cells for both WT and BRM KO together at D4, D6 and D10 of differentiation (**e**) and separately for WT (**f**) and BRM KO cells (**g**). **h-i**, Transcriptional trajectory analysis from single cell data showing step wise transition of WT cell from D4 to D10, and sudden acquisition of neural fate in BRM KO cells (**h**), with cardiac and neural marker expression shown along differentiation trajectory (**i**).

Quantitative gene expression analysis at D4 revealed minimal expression of pluripotency (*Nanog and Sox2*) and paraxial mesoderm markers (*Tbx6, Msgn1*), while primitive streak marker (*T*) and mesoderm precursor (*Pou5f1* or *Oct4*) showed minimal changes between genotypes (Extended Data Fig. 2a), confirming proper cardiogenic mesoderm differentiation of *Brm*^*−/−*^ cells. The absence of *Tbx6*+/*Sox2*+ or *T*+/*Sox2*+ cells precluded the possibility that *Brm*^*−/−*^ cells represent neuromesodermal precursors, especially given that our differentiation lacks retinoic acid^20,21^. At D4, we did not observe transcriptional changes in neuroectodermal markers^22^ between WT and *Brm*^*−/−*^ cells. Indeed, the continuous expression of POU5F1 and absence of SOX2 indicated that D4 cells are derived from a mesendoderm rather than an neuroectoderm lineage^23,24^, suggesting *Brm*^*−/−*^ cells acquire neural lineage after D4 (Fig. 2c, Extended Data Fig. 2a).

BRG1, a paralog of BRM, has important roles in cardiogenesis^5,25,26^. To compare the role of *Brg1* to that of *Brm* in cardiac differentiation, we induced genetic *Brg1* deletion^25,27^ at D2 and analyzed the effects at D4 and D10 by single cell RNA-seq (Extended Data Fig. 3a). *Brg1* loss did not affect D4 transcription broadly (Extended Data Fig. 3b, c). However, at D10, WT and *Brg1* KO cells clustered separately (Extended Data Fig. 3b). Marker analysis revealed that *Brg1*-deficient cells formed very few cardiac myocytes and instead formed endothelial cells, fibroblasts, neural progenitors, and developmentally-arrested progenitors (Extended Data Fig. 3b-e). Unlike *Brm*^*−/−*^ cells, we did not observe TUBB3 staining in *Brg1* KO cells (Extended Data Fig. 3f).

These results show that under directed differentiation conditions, *Brm*^*−/−*^ cells form specified cardiac mesodermal cells, however then undergo a radical fate change towards a constellation of non-cardiac, largely neuronal cell types. Loss of Brg1 also had a similar but less severe phenotype, indicating essential but distinct roles of BAF complex ATPases in regulating lineages during differentiation^28,29^.

### BRM modulates dynamic chromatin accessibility

We next used ATAC-seq^30^ to examine BRM’s role in modulating chromatin accessibility during cardiac differentiation (Fig. 3a). Although gene expression was minimally affected at D4, we found significant changes (FDR<0.05, fold change >2) at 3320 chromatin regions, 98.3% of which showed reduced accessibility upon *Brm* loss. These sites were enriched for genes involved in cardiac and other developmental pathways (Fig. 3b). At D6 and D10 8814 and 5391 regions were significantly changed (FDR<0.05, fold change >2), respectively, between genotypes (Supplementary Sheet 3). Consistent with the RNA-seq data, at D6 the differentially closed chromatin in *Brm*^*−/−*^ cells was near regulatory elements of cardiac development genes, including TFs such as *Gata4, Tbx5, Nkx2-5, Myocd, Hand2, Mef2c*, and cardiac functional genes such as *Ttn, Myl2, Myh6, Myh7, Actc1.* In contrast, D6 *Brm*^*−/−*^ cells had newly open chromatin near genes involved in non-cardiac differentiation processes, including some neural genes (e.g. *Pax6*) (Fig. 3c), consistent with initiation of neural gene expression. By D10, a clear and strong association was found with accessible chromatin near neural differentiation-related genes (Fig. 3d). Comparison of temporal accessibility patterns for select genes revealed that BRM maintains accessibility near cardiac genes throughout differentiation beginning at D4 (Fig. 3e, Extended Data Fig. 4a). In contrast, loss of BRM induced or maintained accessibility near neural genes at or after D6 (Fig. 3e, Extended Data Fig. 4b).

**Fig. 3.**
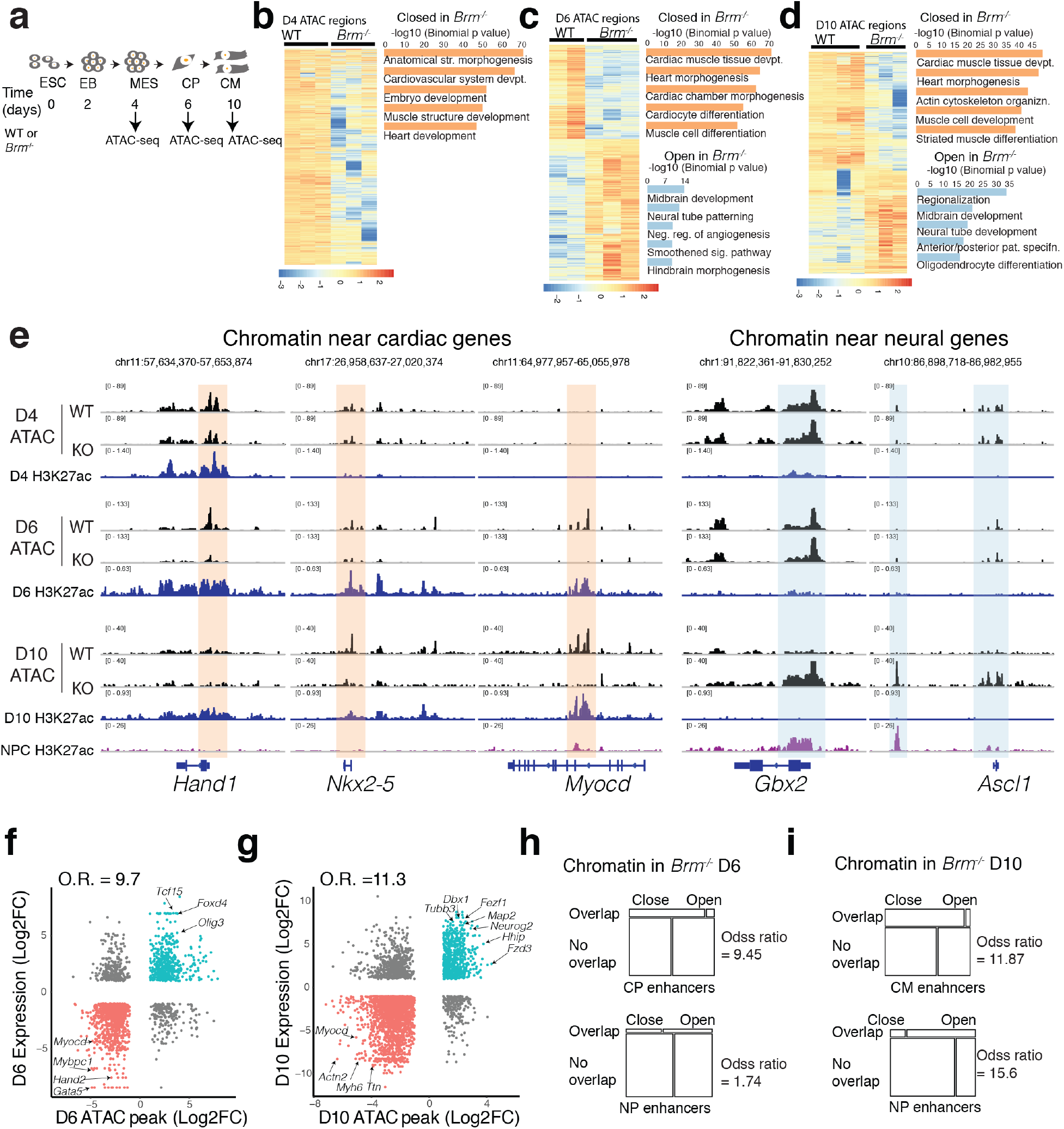
BRM remodels enhancer chromatin to open cardiac and close non-cardiac regulatory regions. **a**, Schematics of cardiac differentiation and timing of ATAC-seq**. b-d**, Heat map of significantly altered ATAC-seq peaks in WT and BRM KO at D4 (**b**), D6 (**c**) and D10 (**d**). GREAT enrichment (two nearest genes within 100Kb) of gene ontology (GO) biological processes shown on the right. **e**, Example browser tracks show ATAC-seq regions over promoter and regulatory regions of key cardiac and neural genes along with D4, D6, D10 and neural progenitor cell enhancer H3K27ac tracks. **f-g**, ATAC-seq peak strengths are correlated with the neighboring BRM regulated genes (within 100Kb, FDR<0.05, ±2 fold) at D6 (**f**) and D10 (**g**). **h-i**, BRM-mediated open and close chromatin regions compared with cardiac and neural progenitor enhancers. Closed and open chromatin in *Brm^−/−^* at D6 (**h**) and at D10(**i**) are compared with respective cardiac and neural progenitor enhancers.

Altered chromatin accessibility was highly correlated with gene expression changes at D6 and D10 (Figs. 3f and 3g). Comparing BRM-mediated chromatin accessibility to active enhancers identified by H3K27ac marks^31,32^ showed that BRM promoted accessible regions at CP and CM enhancers (Figs. 3h and 3i). Conversely, BRM-mediated closed chromatin associated significantly with neural progenitor enhancers in CMs^32^. Motifs enriched in ATAC-seq peaks that were significantly depleted in *Brm*^*−/−*^ included those for cardiac-related transcription factors (GATA4, MEF2C, HAND2) while peaks newly opened in absence of BRM were enriched for motifs for neuronal TFs (SOX2, OCT6/8 (POU3F1/3), OTX2, LHX2/3, RFX; Extended Data Fig. 4c). Thus, BRM promotes open chromatin at cardiogenic genes, while subsequently establishing or maintaining the inaccessible state of non-cardiac (including neural) enhancers.

### Timing of BRM function

To more precisely pinpoint the timing of BRM function, we created an auxin-inducible degron ES cell line^33^ to rapidly deplete BRM (Extended Fig. 4d). Continuous auxin application largely recapitulated the impaired cardiac differentiation of *Brm*^*−/−*^ cells, with a low level of remaining TNNT2+ cells likely due to incomplete BRM depletion in a proportion of cells (Extended Data Fig. 4e)^33^. Depletion of BRM prior to D4 impaired differentiation, whereas subsequent depletion did not greatly affect global cardiac differentiation (Extended Data Fig. 4f). BRM loss did not affect ESC pluripotency or self-renewal (Extended Data Fig. 5a). These results indicate a critical role for BRM after exit from pluripotency, and during cardiac mesoderm formation, but before mesoderm to CP transition.

**Fig. 4.**
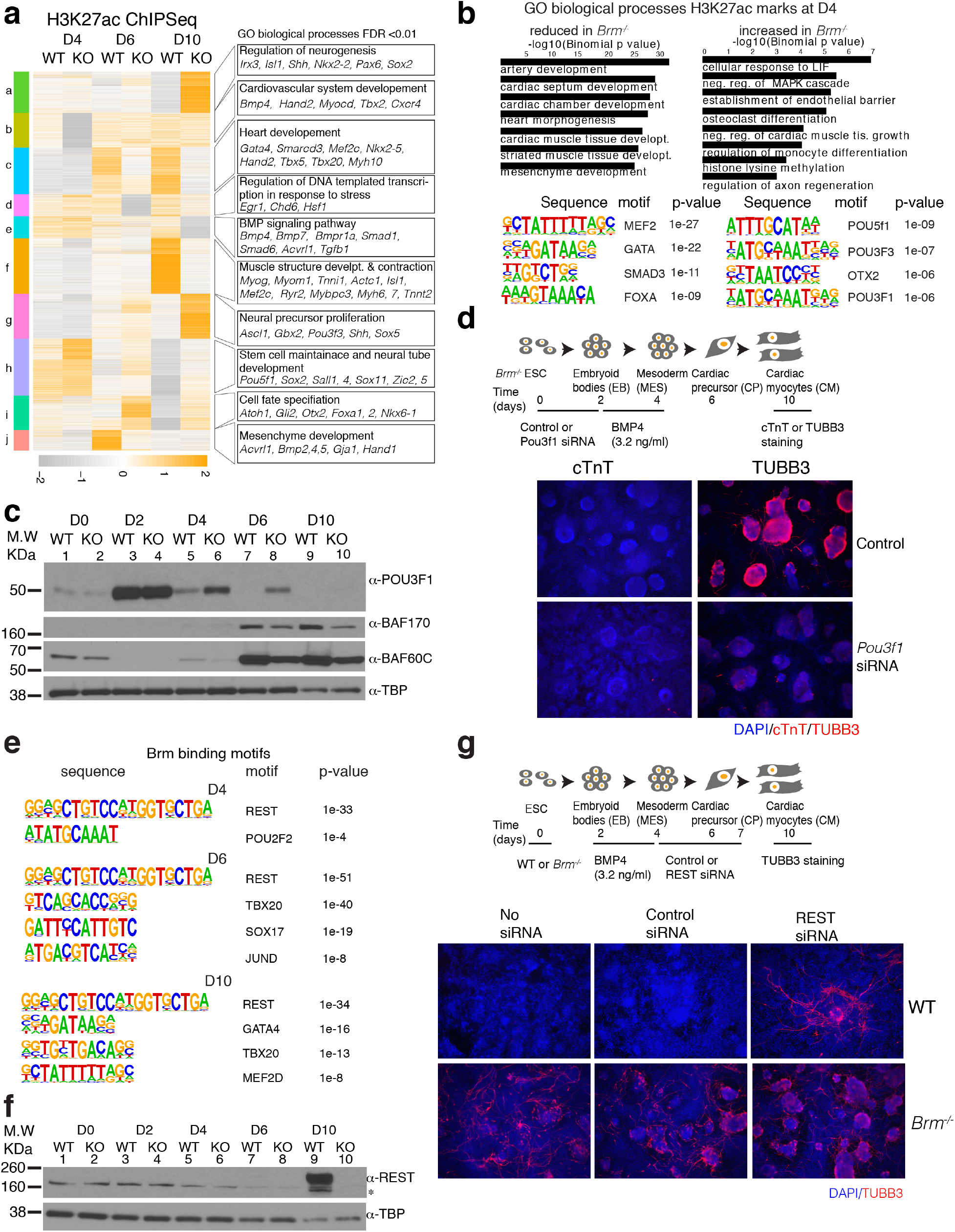
BRM modulates expression of POU3F1 and REST to repress neurogenesis and facilitate cardiogenesis. **a**, Heat map of significantly affected (FDR<0.05, fold change 2) H3K27ac peaks due to loss of BRM at D4, D6 and D10 of differentiation. GREAT analysis of GO biological processes enriched (within 1mb) are shown to the right of the clusters. **b**, Biological process enriched at D4 for sites that reduced (left) or gained (right) H3K27ac in absence of BRM. Corresponding motifs enriched are shown underneath. **c**, Western blot of indicated proteins in WT and BRM KO cells during cardiac differentiation. **d**, Scheme of *Pou3f1* knockdown during cardiac differentiation followed by immunostaining with cTnT and TUBB3 at D10. **e**, Motifs enriched on BRM binding sites from BRM3xFLAG ChIP-seq peaks at D4, D6 and D10. **f**, Western blot showing loss of REST expression in BRM KO cells. **g**, Scheme of REST knockdown during cardiac differentiation followed by TUBB3 immunostaining at D10.

### Epigenetic regulation of chromatin by BRM

BAF complex subunits been implicated in modulation of chromatin in lineage-specific enhancers^25,34^. BRM specifically facilitates acetylation of H3K27 residues at H3K27me3 enriched Polycomb targets^35,36^. To understand if BRM modulates enhancer landscape via histone modifications, we profiled the effect of BRM loss on H3K27ac and H3K27me marks. At D4 of cardiac differentiation, very few regions were differentially enriched with H3K27me3. At later stages, regions near cardiovascular genes gained H3K27me3 marks upon BRM loss, while PcG-repressed genes involved in early embryo development lost their H3K27me3 marks (Extended Data Fig. 6a).

Conversely, in D6 and D10 *Brm*^*−/−*^ cells, H3K27ac was reduced near genes associated with cardiac muscle development and contraction (Fig. 4a, clusters *c* and *f*, Extended Data Fig. 5c,d, 6c,d), and increased near genes involved in cell fate specification, neurogenesis and regulation of neuron differentiation (Fig. 4a, clusters *a*, *g* and *I*, Extended Data Fig. 5c, d, 6c,d), concordant with ATACseq data. Sites reduced in *Brm*^*−/−*^ cells were enriched for cardiac TFs, while sites that gained H3K27ac marks were enriched for neural TF motifs (Extended Data Fig. 5f, g). At D4, sites with reduced H3K27ac in *Brm*^*−/−*^ cells were enriched near cardiovascular development genes (Fig. 4a, cluster b, Extended Data Fig 5b, e), and had motifs for cardiac TFs. Despite the absence of accessibility changes we observed sites that gained H3K27ac in D4 *Brm*^*−/−*^ cells, including genes involved in stem cell maintenance and neurogenesis (Fig. 4a, cluster *h*), which were enriched for POU or OCT motifs (Fig. 4b).

The enrichment of POU motifs suggested a potential involvement of these TFs in neural induction in *Brm*^*−/−*^ cells. Bulk RNAseq with lower statistical cutoff (raw p-value <0.05) showed a modest increase in *Pou3f1* (*Oct6*) mRNA in *Brm*^*−/−*^ D4 cells; other POU factors were not detected. POU3F1 promotes neural fate by activating neural lineage genes and inhibiting BMP4-dependent transcription^37^. We found that POU3F1 protein was expressed at D2, and thereafter reduced during WT differentiation (Fig. 4c). Despite the low level of mRNA induction, POU3F1 protein was robustly increased at D4 and D6 in *Brm*^*−/−*^ cells (Fig. 4c), suggesting prolonged POU3F1 may initiate the neurogenic gene expression program. Indeed, knockdown of *Pou3f1* in *Brm*^*−/−*^ cells resulted in fewer TUBB3^+^ neuronal progenitor cells and absence of filamentous extensions (Fig. 4d).

To delineate BRM occupancy, we performed chromatin immunoprecipitation followed by sequencing (ChIP-seq) using anti-FLAG antibody on a BRM-3xFLAG tagged strain at D4, D6 and D10 (Extended Data Fig. 7a). BRM bound to 110 regions at D4, 521 regions at D6 and 1188 regions at D10 (Extended Data Fig. 7b-d and Supplementary Sheet 4), consistent with its increasing expression pattern during cardiac differentiation. BRM-bound regions were enriched for genes involved in transcriptional and post-transcriptional regulation of gene expression at D4 and D6, and for regulation of muscle development at D10 (Extended Data Fig. 7e). Motif analysis revealed enrichment of REST motifs at all stages, along with cardiac TFs at D10 (Fig. 4e). Moreover, BRM deletion abrogated REST expression at D10 (Fig. 4f). REST knockdown from D4 to D7 resulted in ectopic expression of TUBB3^+^ cells at D10 (Fig. 4g), suggesting BRM controls expression of REST to specifically represses neural lineage genes during cardiac differentiation.

### Signal-dependent rescue of anomalous differentiation

The BRM degron experiments suggested that BRM is most critical during cardiac mesoderm induction (Extended Data Fig. 4f). At this stage, BRM regulates neuronal lineage inducing factor POU3F1, which can counteract BMP signaling^37^. BMP4 concentration at this stage is finely regulated to ensure proper cardiac differentiation ^38,39^. In our system, supraphysiological exogenous BMP4 concentration inhibited cardiac differentiation of WT ESCs but rescued cardiac differentiation of *Brm*^*−/−*^ ESCs (Fig 5a, Extended Data Fig. 8a). High BMP4 also repressed prolonged POU3F1 expression in *Brm*^*−/−*^ cells and normalized expression of BAF60c and REST (Fig. 5b). Loss of BRG1, however was not similarly compensable by increasing BMP4 concentrations (Extended Data Fig 8b).

**Fig. 5.**
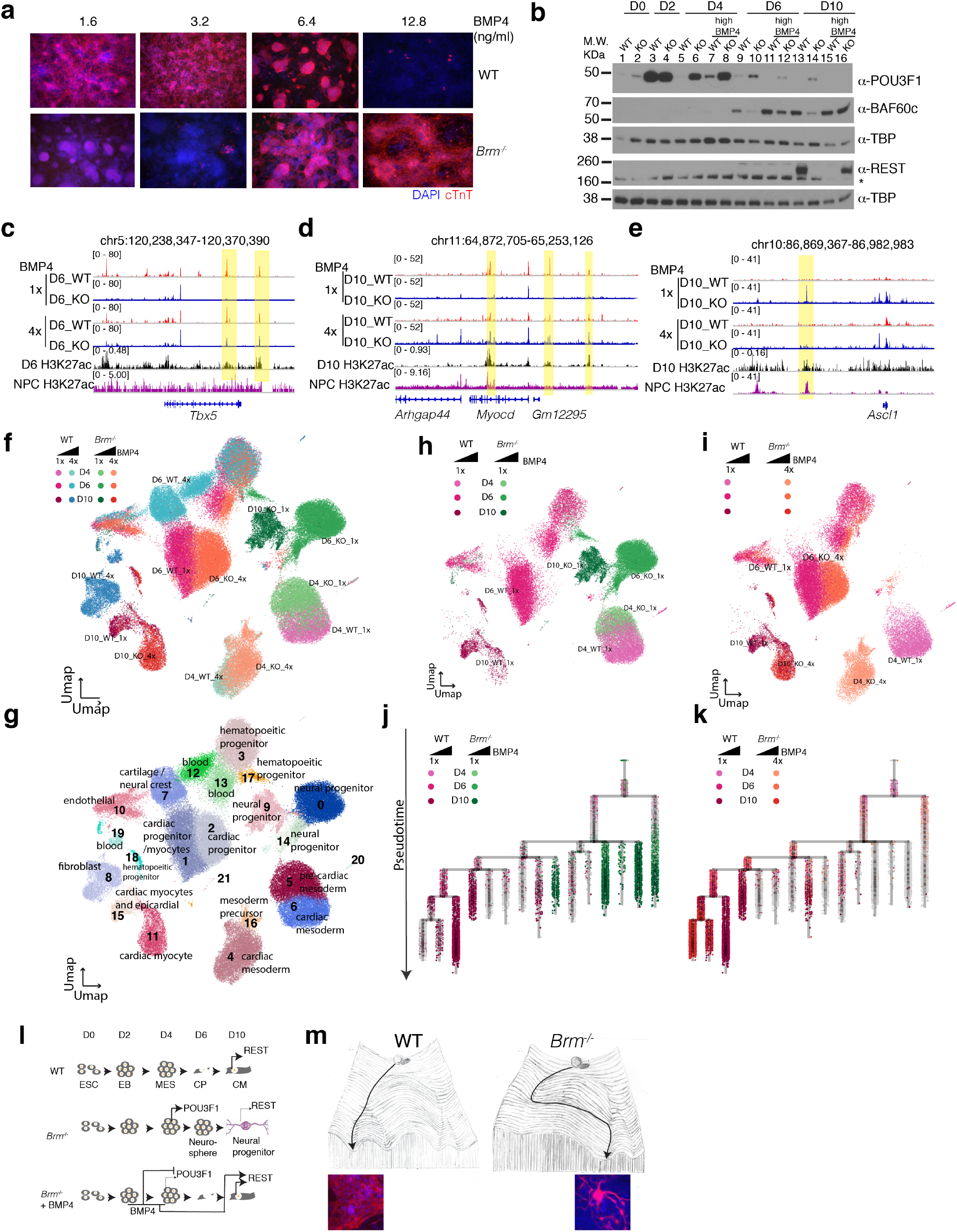
BMP4 signaling rescues loss of BRM phenotype. **a**, Immunostaining of WT and BRM KO cells in presence of increasing concentrations of exogenous BMP4. BMP4 treatment occurred at D2 to D4 of differentiation. **b**, Western blot showing repression of POU3f1 and re-expression of REST and BAF60c in presence of high BMP4 in BRM KO cells. **c-e**, Browser tracks showing re-opening of closed enhancer regions in BRM KO cells in presence of high BMP4 at *Tbx5* (**c**) and *Myocd* (**d**) and closing of open chromatin at *Ascl1* (**e**) loci at the yellow highlighted regions. **f-g**, Single cell RNAseq data projected on a UMAP space showing both WT and BRM KO genotypes (**f**), clusters with inferred cell types (**g**) at D4, D6 and D10 of differentiation induced with normal (1x, 3.2 ng/ml) and high (4x, 12.8ng/ml) BMP4 concentrations. **h-i**, Both WT and BRM KO at normal BMP4 concentration show divergence in clustering in the UMAP space (**h**), while WT at normal BMP4 shows similar clustering with BRM KO at high BMP4 concentrations and occupy the same UMAP space (**i**). **j-k**, Transcriptional trajectory analysis showing divergence in differentiation path for WT and BRM KO cells at normal BMP4 concentrations (**j**), while WT at normal and BRM KO at high BMP4 concentration make step-wise transition to form cardiomyocytes (**k**). **l**, Model showing at normal BMP4 concentration, WT cells form cardiomyocytes and express neuronal inhibitor REST. In Brm^−/−^ cells, neurogenic factor POU3F1 fails to be repressed whereas REST is repressed, resulting in neural progenitor formation. Brm^−/−^ cells induced with high BMP4 repress POU3F1 and re-express REST, forming cardiac myocytes. **m**, Waddington landscape depicting WT differentiation forming cardiomyocytes, while Brm^−/−^ cells continue on a cardiac differentiation path before undertaking a saddle-node bifurcation to form neural progenitor cells.

BMP4-mediated BRM rescue restored accessibility of cardiac enhancers (Fig. 5c-e, Extended Data Fig. 8c-e, see boxed regions and 8f). Conversely, neural progenitor enhancers were inaccessible in *Brm*^*−/−*^ cells with high BMP4 (Fig. 5e, Extended Data Fig. 8g).

Single cell RNAseq analyses revealed BMP4-dependent changes in gene expression. At D4, WT and *Brm*^*−/−*^ cells differentiated with normal BMP4 concentrations (3.2 ng/μl) clustered together in the same UMAP space, but separated from the high BMP4 (12.8 ng/μl) samples (Fig. 5f). Later in differentiation, high BMP4 WT cells clustered separately from normal BMP4-treated cells, forming endothelial and hematopoietic progenitors, and eventually fibroblasts and blood cells (Fig. 5f,g). As expected, *Brm*^*−/−*^ cells at normal BMP4 formed mostly neural progenitor clusters (Fig. 5h) separate from WT cells. In contrast, *Brm*^*−/−*^ cells at high BMP4 clustered with WT cells treated with normal BMP4 at D6 and D10, and expressed CP and CM markers (Figs. 5f, i, Extended Data Fig. 9a and Supplementary Sheet 5).

URD trajectory analysis with *Pou5f1+* expressing clusters as root and D10 clusters as tips placed WT and *Brm*^*−/−*^ cells at normal and high BMP4 concentrations at the trajectory root. At normal BMP4, WT cells followed a step-wise trajectory forming cardiomyocytes, while *Brm*^*−/−*^ cells followed a different path (Fig. 5j), however *Brm*^*−/−*^ cells at high BMP4 formed cardiac progenitors and cardiomyocytes by following an almost identical trajectory to WT cells at normal BMP4 (Fig. 5k, Extended Data Fig. 9b). Thus, simply modulating BMP4 signaling compensated for the absence of BRM and completely restored the normal path of cardiac differentiation.

How BMP4 might change differentiation path of *Brm*^*−/−*^ cells is not clear. We confirmed that loss of BRM simply did not change BMP4 availability to the cells (Extended Data Fig. 9c). BMP-dependent gene regulatory networks are highly robust, and self-regulate in part by modulating transcriptional noise^40,41^. To evaluate if gene expression noise is regulated by BRM or BMP4, we calculated cell-to-cell gene expression variability^42^ and single cell entropy^43^ to predict differentiation potential. At normal BMP4 both WT and *Brm*^*−/−*^ cells had similar dispersion and entropy metrics, both of which increased in presence of high BMP4 at D4 (Extended Data Fig 9c and 9d). This suggests increased intrinsic gene expression noise could participate in BMP4-dependent modulation of the *Brm*^*−/−*^ transcriptional state.

### In vivo requirement for BRM

*Brm*^*−/−*^ mice are viable, although the nature of the original allele has been questioned^6,44^. Our independent mouse line with an 8bp deletion at Exon 2 resulting a premature stop codon and loss of BRM protein produced pups at Mendelian ratios (Extended Data Figs. 10a-c) confirming viability of *Brm* knockout. Viability may be due to compensatory overexpression of BRG1 in *Brm* knockout tissues (Extended Data Fig. 10d), which did not occur in our directed differentiation system (Extended Data Fig. 10e).

## Discussion

Along the “landscape” of cell fate decisions, epigenetic regulators are key determinants of transition states. This is apparent in cancer, where new attractor states are formed that result in anomalous differentiation or dedifferentiation. In normal development however, only scant examples exist of natural transdifferentiation^21,45,46^, pointing to stability and robustness of canalization in vertebrate differentiation. Here we show that this stability during cardiac differentiation requires safeguarding by BRM.

Artificially forced reprogramming overcomes cell states^47^, and certain chromatin remodeling factors including BRM are important safeguards against reprogramming^48–51^. Conversely, other BAF complex subunits (e.g. BRG1, BAF60c) enhance reprogramming^52–54^. Reprogramming of fibroblasts to neurons involves transient competition between myogenic and neural gene expression programs, evidence that genome plasticity can transcend germ layer specification^55^. It is likely that in the absence of BRM, deregulation of neurogenic TFs, e.g. POU3F1, activates a cascade of neural gene expression in the context of broadly deregulated enhancer accessibility (Fig. 5l). That we observe this “self-reprogramming” in a directed differentiation context, but not in the complete organism indicates that the cues provided in vitro are strictly narrow parameters, while in vivo they are likely highly buffered. Indeed, increased BRG1 in *Brm*^*−/−*^ mice and rescue of *Brm*^*−/−*^ cells by elevated BMP signaling, indicates that loss of BRM is compensable.

Our findings indicate that BRM maintains developmental canalization of committed mesodermal precursors by providing an epigenomic state that favors a limited range of transition states, and that in its absence an unstable state induces a transition akin to a saddle node bifurcation (Fig. 5m). We highlight the fragility of the differentiation path, challenging the concept of highly robust developmental canalization, with important implications for understanding the stability of gene regulation in differentiation, and for deregulated gene expression in disease.

## Supporting information

Supplemental Sheet 5

Supplemental Sheet 2

Supplemental Sheet 1

Supplemental Sheet 4

Supplemental Sheet 3

## Methods

### Cell culture and in-vitro differentiations

Mouse embryonic stem cells (ESCs) were cultured in media containing fetal bovine serum (FBS) and leukemia inhibitory factor (LIF) without feeder mouse embryonic fibroblast cells with daily media change at 37°C, 7% CO2 and 85% relative humidity. CMs were differentiated as described previously^31,38^. Briefly, mouse ESCs were cultured in presence of ascorbic acid (50μg/ml) in suspension cultures without LIF and Serum for 2 days to form embryoid bodies (EBs). EBs were dissociated and treated for 2 days with VEGF, Activin A and BMP4 to induce cardiac mesoderm which were subsequently dissociated and cultured as monolayer in presence of FGF-basic (FGF2) and FGF10 for 6 days to form beating cardiac myocytes. *Brg1* was deleted in presence of 200 nM 4-hydroxytamoxifen for 48 h with control cells treated similarly with tetrahydrofuran^5,25,56^. Neural stem cell differentiations were carried out in presence FGF2 and epidermal growth factor (EGF), with growth factor removal forming neuronal progenitor cells as described previously^57^.

### Cell line and mouse line generation

BRM was targeted using CRISPR/Cas9 with sgRNA targeting exon 2 following the described protocol^58^. sgRNA were cloned to a BbsI-digested pX330 vector (Addgene Cat #42230) by annealing the following primers: 5′ **caccg** GTCCACTGTGGATCCATGAA 3′ and 5′ **aaac** TTCATGGATCCACAGTGGAC **c** 3′ (bold indicates the BbsI digestion site). For construction of BRM-3xFLAG tag line, we followed a similar strategy to insert 3xFLAG tag sequence between the stop and penultimate codon using the following primers to clone sgRNA to the BbsI site of pX330 vector : 5’ caccg CTGATAACGAGTGACCATCC 3’ and 5’ aaac GGATGGTCACTCGTTATCAG C 3’. The following sequence was inserted to the upstream homology sequence for the insertion of 3x-FLAG tag: 5’ ggaggcggtggagcc GAC TAC AAG GAC CAC GAC GGC GAC TAC AAG GAC CAC GAC ATC GAC TAC AAG GAC GAC GAC GAC AAG TGA 3’. BRM targeting vectors were constructed by cloning 450 to 500 bp of DNA upstream and downstream of midpoint of sgRNA target site into KpnI-XhoI and BamH1-NotI sites of pFPF (a derivative of Addgene plasmid #22678 in which neomycin is replaced with puromycin cassette). BRM-AID strain was constructed following a previously-described strategy^33^. Briefly, pEN244-CTCF-AID_71-114_-eGFP-FRT-Blast-FRT plasmid (addgene Cat#92140) was digested with BamH1 and Sal1 and replaced the 3’ and 5’ homology of *Ctcf* with that of *Brm* respectively. The following primers were used to clone an sgRNA to pX330 vector: 5’ CAC CCT GAT AAC GAG TGA CCA TCC 3’ and 5’ GAC TAT TGC TCA CTG GTA GGC AAA 3’. 2.5 μg of each of the sgRNA plasmid, plus 20 μg of *Brm* targeting constructs were used for transfection. Single clones were selected, grown, PCR genotyped and DNA sequenced.

For constructing a *Brm* mouse strain, we used CRISPR/Cas9 with the exact same exon 2 sgRNAs as in the cell line cloned to a BbsI-digested pX330 vector by annealing oligos: 5’ **caccg** GTCCACTGTGGATCCATGAA 3’ & 5’ **aaac** TTCATGGATCCACAGTGGAC **c** 3’. In-vitro transcribed RNA and CAS9 protein complex and were injected to the embryos and transferred to 0.5 dpc pseudo-pregnant female mice. We obtained a mouse line with 8bp deletion resulting in a premature stop codon, confirmed by genotyping PCR sequencing and loss of BRM protein by western blot

### siRNA mediated knockdown

RNA knockdown were carried out using Lipofectamine-RNAiMax reagent (ThermoFisher, 13778150) and pre-designed siRNA against POU3F1 (Sigma, SASI_Mm02_00319981) and REST*(Sigma*, `SASI_Mm01_00196017`) mRNAs. Control siRNA were used as negative controls (Sigma, SIC001-10NMOL). Briefly, cells were split, washed and suspended in suspension culture plates (for D0 differentiation) or monolayer (D4 differentiation). siRNAs (3μl of 10μM conc.) and RNAiMax (7 μl) were mixed separately with 75 μl Optimem (Thermofisher, 31985062). Knockdown was initiated by mixing both siRNA and RNAiMAX suspensions together, incubated for 5 mins at RT. The entire 160 μl of silencing mix were added dropwise to 1ml culture or scaled accordingly.

### Nuclear extracts and Western blot

Nuclear extracts were prepared using protocols described previously^59^. Western blotting was performed using standard techniques with PVDF membranes. Primary antibodies used were anti-BRG1 (Abcam, ab110641, 1:1000), anti-FLAG (Sigma, F1804, 1:1000), anti-BAF170 (Bethyl, 1:1000, A301-39A), anti-BAF60c (Cell Signaling Technology, 62265, 1:1000), anti-REST (EMD-Millipore, 07-579, 1:1000), anti-POU3F1 (Abcam, ab126746, 1:1000), or anti-TBP (Abcam, ab51841, 1:2000), Vinculin (Sigma-Aldrich V9131, 1:1000), phospho-Smad (CST 9511, 1:1000) and, Smad1 (CST 9743, 1:1000). Secondary antibodies used were donkey anti-rabbit IRDye 800cw (Licor, 926-32213, 1: 10,000), donkey anti-mouse IRDye 800cw (Licor, 925-32212, 1: 10,000) and donkey anti-goat IRDye 680cw (Licor, 925-68074-1:10,000), HRP-linked-anti-mouse (Cell Signaling Technology, 7076, 1:10000) or HRP-linked-anti-rabbit (Cell Signaling Technology, 7074, 1:10000).

### Immunofluoresence

Cells in monolayer were fixed for 30mins in 4% para-formaldehyde, permealized in 0.1% Triton X and 5% goat serum in PBS for 1 hr and incubated with primary antibody (anti-FLAG (Sigma, F1804, 1:300), anti-OCT4 (R&D, MAB1759, 1:300), anti-SOX2 (Abcam, ab97959, 1:300), anti-NANOG (Abcam, ab80892, 1:300), anti-cardiac Troponin T (*Thermo Scientific*, *MS-295-P*, 1:100), or TUBB3 (BioLegend, 8012 1:5000) overnight. They were then washed thrice with 0.1% triton X in PBS, incubated with secondary antibody (Goat anti-mouse Alexa 594 (Invitrogen, A11005, 1:1000), Goat anti-rabbit Alexa594 (Invitrogen, 110037, 1:1000) or Donkey-anti-goat AlexaFluor594, 1:1000) for 1hr at RT. Wells were washed thrice and stained with DAPI (1:1000 dilution) for 1-2 min followed by a PBS wash. Images were taken in Keyence confocal microscope at 10x or Zeiss Spinning Disk microscope at 63x (for Extended Data Fig. 5a) magnification.

### Flow cytometry

At D10 of differentiation, WT and BRM KO cells were dissociated using TrypLE and fixed with 4% methanol-free formaldehyde. Cells were washed with PBS and permeabilized using FACS buffer (0.5% w/v saponin, 4% Fetal Bovine Serum in PBS). For evaluation of differentiation efficiency, cells were stained with a mouse monoclonal antibody for cardiac isoform Ab-1 Troponin at 1:100 dilution (ThermoFisher Scientific #MS-295-P) or the isotype control antibody (ThermoFisher Scientific #14-4714-82) for 1 hour at room temperature. After washing with FACS buffer, cells were stained with goat anti-mouse IgG Alexa 594 secondary antibody at 1:200 dilution (ThermoFisher Scientific #A-11005) for 1 hour at room temperature. Cells were then washed with FACS buffer, stained with DAPI for 2 minutes, rinsed, and filtered with a 40-micron mesh. At least 10,000 cells were analyzed using the BD FACS AriaII and results were processed using FlowJo (BD Bioscience).

### Bulk RNA-seq

Total RNA was isolated from biologically triplicate samples using miRNeasy micro kit with on-column DNase I digestion (Qiagen). RNA-seq libraries were prepared using the Ovation RNA-seq system v2 kit (NuGEN). Libraries from the SPIA amplified cDNA were made using the Ultralow DR library kit (NuGEN). RNA-seq libraries were analyzed using Bioanalyzer, quantified using KAPA QPCR and paired-end 100 bp reads were sequenced using a HiSeq 2500 instrument (Illumina). RNA reads were aligned with TopHat2^60^, counts per gene calculated using feature Counts^61^ and edgeR^62^ was used for the analysis of differential expression. K-means clustering and pheatmap functions in R were used to cluster and generate heatmaps. GO enrichment analysis were performed using GO Elite^63^.

### Single cell RNA-seq

Single-cell libraries were prepared using Single Cell 3’ Library Kit v2 (10x Genomics) according to the manufacturer’s protocol. Briefly, about 10, 000 cells were suspended in 0.04% ultrapure BSA–PBS (McLab, #UBSA-500) for GEM generation. GEMs were reverse transcribed, and single stranded DNA were isolated and cleaned. Then cDNA was amplified twice, fragmented, end-repaired, A-tailed and index adaptor ligated, with Ampure cleanup (Beckman Coulter) after each step. Libraries were PCR amplified and cleaned with Ampure beads before shallow sequencing in a NextSeq 500. Read depth normalized libraries were re-sequenced in a NovaSeq sequencer (Illumina).

Sequencing reads were aligned using CellRanger 2.0.2 or 3.0 to the mm9 mouse reference genome. cellranger *aggr* was used to generate an aggregated read normalized data matrix of samples. The filtered gene matrix was subsequently used to create a Seurat object for QC and tSNE or UMAP visualizations as described in https://satijalab.org/seurat/tutorial^64^.

### Seurat analysis

Seurat package v2.3.4 was used to analyze single cell RNA sequencing data. Cells were filtered to remove dead cells and doublets. After log-normalization, sources of unwanted variation, including differences in the number of UMI, number of genes, percentage of mitochondrial reads and differences between G2M and S phase scores were regressed using the *ScaleData* function. Clustering was performed using the top 30 principal components and visualized using Uniform Manifold Approximation and Projection (UMAP)^17^.

Differential gene expression tests were run using the *FindMarkers* function with min.pct set to 0.1 and logfc.threshold set to 0.25. Selected differentially expressed genes with an adjusted p-value less than 0.05 from the Wilcoxon Rank Sum test were then displayed using the Dotplot function.

### Cell trajectories and pseudotime analysis

Single Cell Analysis in Python (Scanpy), version 1.4.5, was used for finding highly variable genes (HVGs), computing dimensionality reduction, regressing unwanted sources of variation, and building developmental trajectory. Two thousand HVGs were selected within each differentiation time separately and merged, to capture differentiation-specific genes^64^ Variations were regressed from HVGs that encode for ribosomal and mitochondrial proteins. HVGs were then scaled to unit variance and zero mean. Next, the regressed two thousand HVGs were decomposed to fifty principal components using the SciPy, version 1.41, ARPACK Singular Value Decomposition (SVD) solver. A k-nearest neighbor graph was then constructed from a local neighborhood size of ten, using thirty principal components (PCs), the euclidean distance metric, and the connectivity estimation of the manifold set to Unified Manifold Approximation Projection (UMAP) The louvain-graph based clustering algorithm was then run at a resolution of 1.0 on the k-nearest neighbor graph^65^. A developmental trajectory was resolved by assessing the connectivities of the louvain clusters from the k-nearest neighbor graph, using partition-based graph abstraction (PAGA)^18^. Finally, the UMAP embedding was recomputed using the PAGA initialization to visualize the developmental trajectory at single cell resolution.

Pseudotime analysis was performed using the URD package^19^ (version 1.0.2). A single expression matrix with data from three timepoints and WT and *Brm*^*−/−*^ in low and high BMP4 conditions was processed in Seurat v2.3.4, as described above. The object was then down-sampled to retain 5000 cells per sample in the low BMP4 dataset or 3000 cells per sample in the combined low and high BMP4 dataset. The down-sampled object was converted to an URD object using the *seuratToURD* function. Cell-to-cell transition probabilities were calculated by setting the number of nearest neighbors (knn) to the square root of total cells in the object. *POU5F1+* clusters from day 4 were set as ‘root’ and all day 10 clusters were set as ‘tip’ cells. An URD tree was constructed by simulating biased random walks from each tip cluster to root.

### Signaling Entropy Analysis

Gene-barcode matrices from single-cell RNA-sequencing of day 4 and 6 differentiation samples were first filtered and normalized using the Seurat package implemented in R. The “LogNormalize” method with a default scaling factor of 10,000 was applied for normalization. Differentiation potency was next estimated for each cell within the datasets using the SCENT algorithm implemented in R, which integrates a cell’s transcriptomic profile with existing protein-protein interaction (PPI) maps to quantify signaling entropy^43^. Higher entropy is an indication of greater developmental potency. A human PPI map compiled from Pathway Commons was used as input for an adjacency matrix (https://github.com/aet21/SCENT). Mouse Ensembl IDs were converted into their human homologues using the *AnnotationTools* Bioconductor package. The resulting set of genes were then integrated with the human PPI network. The entropy value for each cell was normalized to the largest eigenvalue (maximum possible entropy) of the adjacency matrix. Distributions of normalized entropy values for each sample were then plotted for comparison.

### Differential Variability Testing with BASiCS

To assess changes in gene expression variability while accounting for artefactual technical noise and the confounding relationship between variance and mean, single-cell RNA-seq datasets were analyzed via the BASiCS framework as implemented in R^42^. This approach produces gene-specific estimates of residual over-dispersion: a metric describing how greatly a gene’s variability departs from what is expected given its mean expression. Quality control and filtering of gene-barcode matrices was performed using the BASiCS_Filter function with default parameters. Posterior estimates of mean and residual over-dispersion for each gene were computed using a Markov chain Monte Carlo (MCMC) simulation with 40,000 iterations, log-normal prior and regression analysis.

### ATACseq

Assay for transposase-accessible chromatin using sequencing (ATAC-seq) was performed as described^30^ in two to four biological replicates. Briefly, 50,000 cells (>95% viability) were lysed, washed and tagmented for 45 mins and 3 h for CP and CM cells, respectively. DNA was purified and amplified using universal Ad1 and barcoded reverse primers^30^. Libraries were purified, quantified and analyzed on a bioanalyzer and sequenced on a NEB NextSeq 550 sequencer using Illumina NextSeq 500/550 High Output v2 kit (150 cycles). Sequencing image files were de-multiplexed and fastq files generated. Reads (paired end 75 bp) were trimmed and aligned to mouse genome mm9 assembly using Bowtie 2^66^ with a minimum mapping quality score of 30.

Statistically enriched bins with a P-value threshold set to 1×106 were used to call peaks^67^. UCSC genome browser and IGV were used to view the browser tracks. Deeptools package in Galaxy^68^ (usegalaxy.org) was used to pool multiple replicates to generate 1x genome coverage (average of multiple samples) browser tracks. GREAT^69^ was used to generate gene lists near ATACseq sites within 100Kb.

### ChIPseq

Chromatin immunoprecipitations of histone modifications (H3K27ac and H3K27me3) were performed as described^70^ with modifications. Briefly, cells were crosslinked with 1% formaldehyde, and quenched with 0.125 M glycine. Frozen pellets (1 ×107) were thawed, washed, dounced and digested with MNase. Chromatin was sonicated at output 4 for 30s twice with a 1 min pause between cycles then centrifuged at 10,000 g for 10 min at 4°C and stored at −80°C. Chromatin was diluted to fivefold, pre-cleared for 2 h followed by immunoprecipitation with primary antibodies for 12-16 hours at 4°C (H3K27ac, Active motif 39133; H3K27me3, CST 9733s). 5% of samples were used as input DNA. Antibody-bound protein-DNA complexes were immunoprecipitated using 25 μl of M-280 goat anti-rabbit IgG or anti-mouse IgG dyna beads for 2 h, washed a total of ten times with buffers [twice with IP wash buffer 1 containing 50 mM Tris.Cl (pH 7.4), 150 mM NaCl, 1% NP-40, 0.25% sodium deoxycholate and 1 mM EDTA), five times with IP wash buffer 2 containing 100 mM Tris.Cl (pH 9.0), 500 mM LiCl, 1% NP-40 and 1% sodium deoxcholate, and then thrice with IP wash buffer 2 along with 150 mM NaCl for increasing stringency and eluted with 200 μl of elution buffer [10 mM TrisCl (pH 7.5), 1 mM EDTA and 1%SDS) at 65°C for 30 mins. Samples were reverse crosslinked, digested with proteinase K and RNAse A, and purified using AMPure XP beads (Beckman Coulter). To prepare libraries for sequencing, DNA was end repaired, A-tailed, adapter ligated (Illumina TrueSeq) and PCR amplified for 14 cycles. PCR-amplified libraries were size selected (200 – 500 bps)and ampure purified. The concentration and size of eluted libraries was measured (Qubit and Bioanalyzer) before single-end 75bp sequencing using a NEBNextSeq sequencer.

Chromatin IP with anti-FLAG antibodies (Sigma, F1806) to probe for BRM binding sites were performed similarly except following modifications. 1) Cells were double crosslinked with 2 mM disuccinimidyl glutarate (DSG) and 1% formaldehyde. 2) MNase digestion conditions were adjusted to have optimal chromatin digestion yielding fragments sizes of 400 to 1Kb. 3) Chromatin binding to antibody and initial two washes contained either 0.05% (low SDS) or 0.2% (high SDS) conditions. 4) Bound protein was competitively eluted with 0.1mg/ml FLAG peptides (ELIM biopharma) and remaining material at 65°C. We observed better ChIP signal over noise at high SDS samples eluted with the FLAG peptides.

Reads (single end 75 bp) were processed as in ATACseq analysis. The HOMER^71^ motif enrichment package was used to enrich DNA motifs in both ATACseq and ChIP-binding sites. HOMER calculates the q-value of known motifs to statistically confirm to Benjamini-Hochberg multiple hypothesis testing corrections.

### ATAC-seq and ChIP-seq analysis

The raw sequence data in fastq files were aligned to the mouse genome build mm9 using bowtie2 aligner ^66^. Open chromatin regions and regions marked by H3K27Ac for each sample were called using the narrowPeak output of the MACS2 peak caller ^72^. Regions marked by H3K27me3 were called using the BCP ^72^ peak caller. A consensus set of peaks across replicates (across samples for each of the ATAC-seq and the histone modification ChIPs) is defined using the *-everything* followed by *-merge* options of the bedops program ^73^. A peak is included in the consensus set of peaks (for the ATAC-seq data or the particular histone modification ChIP-seq data) if it includes a peak called by the relevant peak caller for at least one of the associated replicates. The number of reads mapping to each of the consensus regions for each of replicates using the *subread featureCounts* program ^61^. This creates a matrix of raw counts - the number of rows equals the number of consensus regions and the number of columns equals the number of samples. Regions that don’t have at least 5 reads in at least 2 of the samples are filtered out. The raw counts matrix is then normalized using edgeR bioconductor ^62,73^ R package. For each data set, a linear model is fit for the mean normalized signal in each of the filtered consensus region. This model allows for the main effects of genotype (BRM KO versus Wild type), differentiation time (D0, D4, D6 and D10), conditions (normalBMP4 vs high BMP4) and the interaction between these two variables. The significance of the regions associated with genotype, condition and/or differentiation time is estimated by testing the combined null hypothesis that the main effects of genotype, differentiation time and the interaction effect of between these two variables are all equal to zero. The heatmap of significantly associated regions (FDR < 0.1) is done using the pheatmap package in R.

## Data availability

Bulk and single cell RNAseq, ATACseq, and ChIPseq datasets have been deposited in GEO under accession number GSE150186.

## Code availability

Codes used to analyze single cell data on Seurat, generate heat maps, UMAPs and URD figures will be available upon request.

## Author Contribution

Project design and direction: B.G.B. and S.K.H. ES cell engineering, in vitro differentiation, gene expression, scRNAseq, ATACseq, ChIPseq and data analysis: S.K.H. Additional scRNA-seq analysis: A.P.B, K.R., under direction of B.G.B and I.S.K. Cell culture: A.M.B., K.S. Data analysis: R.V.D. under direction of L.S.W. Manuscript writing: S.K.H and B.G.B with contribution from all authors.

## Acknowledgements

We thank Natasha Carli, Y. Hao, M. Bernardi, and J. McGuire (Gladstone Genomics Core) for RNA-Seq and 10X Genomics library preparation, the UCSF Center for Applied Technologies for sequencing, Elphege Nora for help with *Brm-AID* strain construction, Vasumathi Kameswaran for help with BRM ChIP-seq, Reuben Thomas for ATAC-seq and ChIP-seq data analysis, J. Zhang (Gladstone Transgenic Core) for knockout mouse production, and K. Claiborn for editorial assistance. This work was supported by grants from the NIH/NHLBI (P01HL089707, Bench to Bassinet Program UM1HL098179, and R01HL114948), to B.G.B; and postdoctoral fellowships from the American Heart Association (13POST17290043), Tobacco Related Disease Research Program (22FT-0079) and NIH training grant (2T32-HL007731 26) to S.K.H. I.S.K. was supported by funds from the Society for Pediatric Anesthesia, Hellman Family Fund, UCSF REAC Award and the UCSF Department of Anesthesia and Perioperative Care. This work was also supported by an NIH/NCRR grant (C06 RR018928) to the J. David Gladstone Institutes and by The Younger Family Fund (B.G.B.).

---

**Extended Data Fig. 1.**
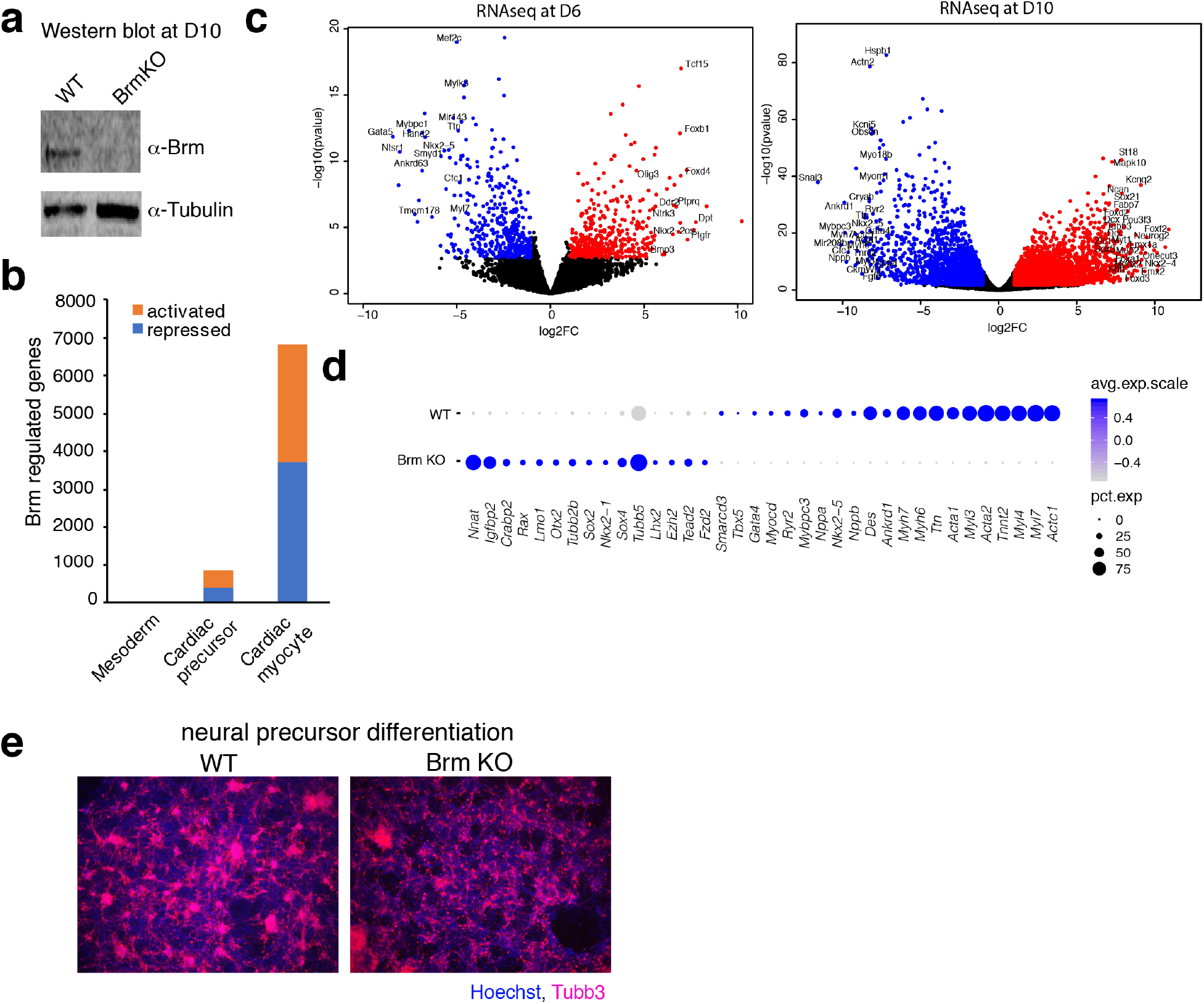
Loss of BRM leads to expression of neural genes in cardiac differentiation and has minimal effect in neural differentiation. **a**, Western blot at D10 of cardiac differentiation of WT and BRM KO cells**. b**, Volcano plots of RNA-seq data showing significantly (FDR<0.05 and fold change > 2) downregulated (blue) and upregulated (red) genes at D6 and D10 stages of differentiation. **c**, Quantification of significantly (FDR<0.05 and fold change > 2) de-regulated genes at D4 (mesoderm), D6 (cardiac precursor) and D10 (cardiomyocyte) stages of differentiation. **d**, Dots plots showing expression of indicated genes from D10 WT and BRM KO single cell RNA-seq data. **e**, TUBB3 immunostaining of WT and BRM KO cells differentiated to neural precursor cells.

**Extended Data Fig. 2.**
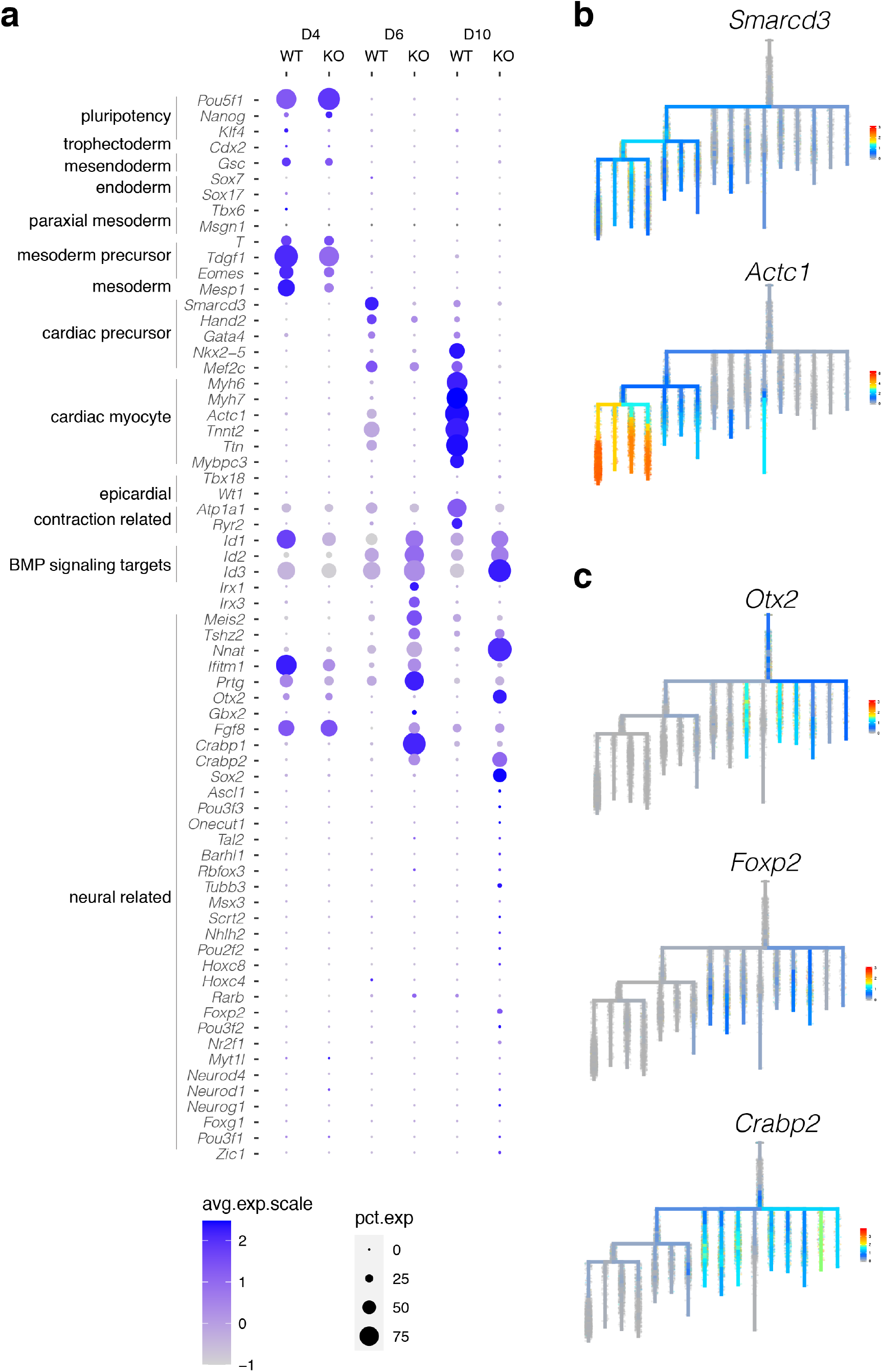
Brm loss leads to expression of neural genes after D4 of differentiation. **a**, Dot plots showing quantitative bulk changes in gene expression between WT and BRM KO cells at D4, D6 and D10 stages of differentiation for early developmental, cardiac mesoderm, precursors, myocytes and genes enriched in BRM KO cells and a select set of genes involved in neuroectoderm development. **b-c**, Feature plots of developmental trajectory analysis using URD for selected cardiac (**b**) and neural genes (**c**)

**Extended Data Fig. 3.**
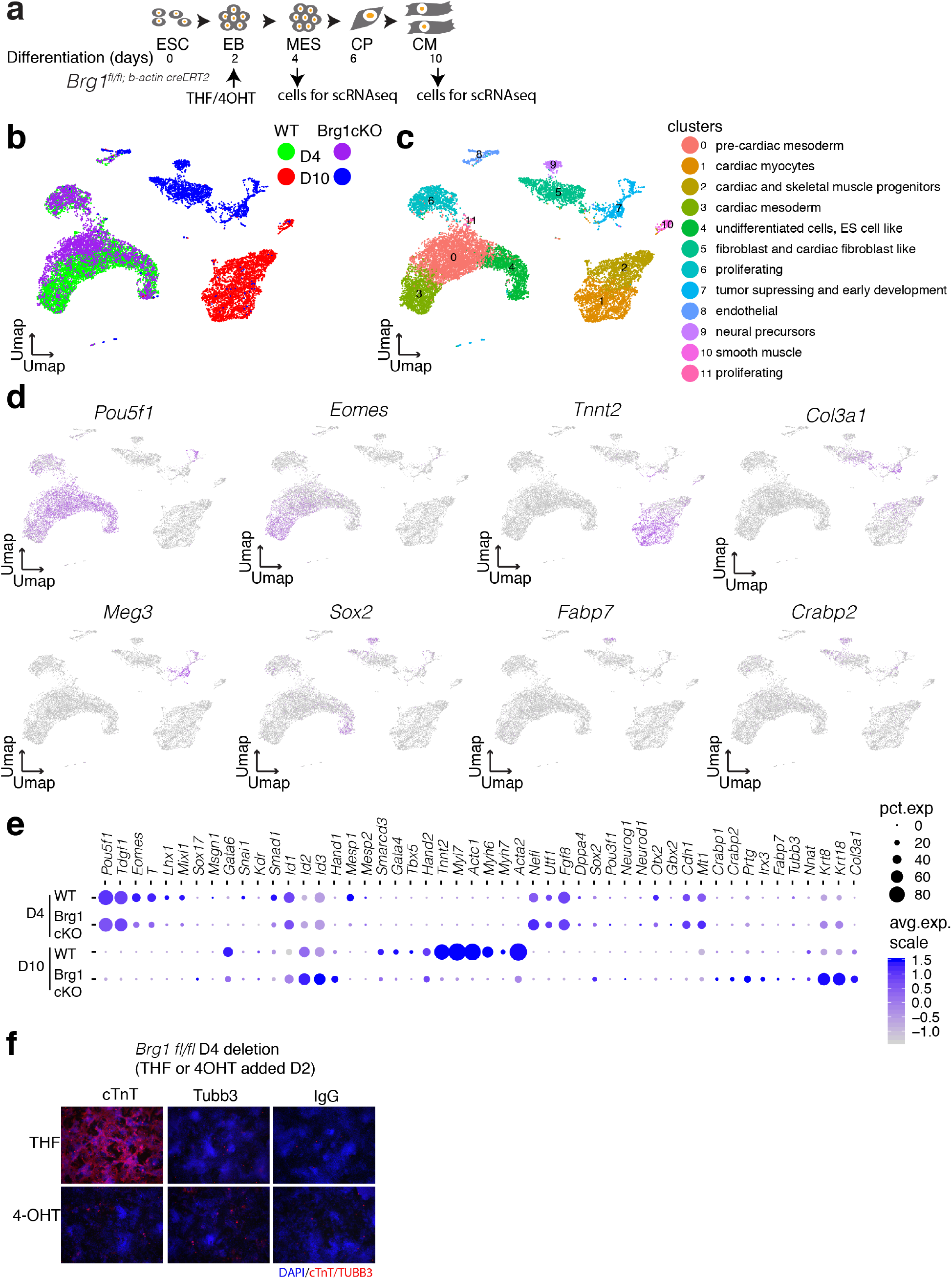
Loss of BRG1 early in differentiation leads to formation of non-cardiac cell types. **a**, Scheme of cardiac differentiation showing timing of induction with 4-hydroxy tamoxifen or the control tetrahydrofuran and scRNA-seq. **b-d**, UMAPs of single cell RNA-seq data at D4 and D10 of differentiation of WT and conditional BRG1 KO genotypes (**b**), clusters with inferred cell types (**c**) and feature plots of expression of indicated genes (**d**). Dot plots comparing gene expression quantification of WT and conditional BRG1 KO at D4 and D10 of differentiation. **e**, Cardiac troponin T and TUBB3 immunostaining at D10 for WT and BRG1 cKO cells deleted at D2 of differentiation. **f**, Immunostaining with cTnT at D10 with increasing concentration of BMP4.

**Extended Data Fig. 4.**
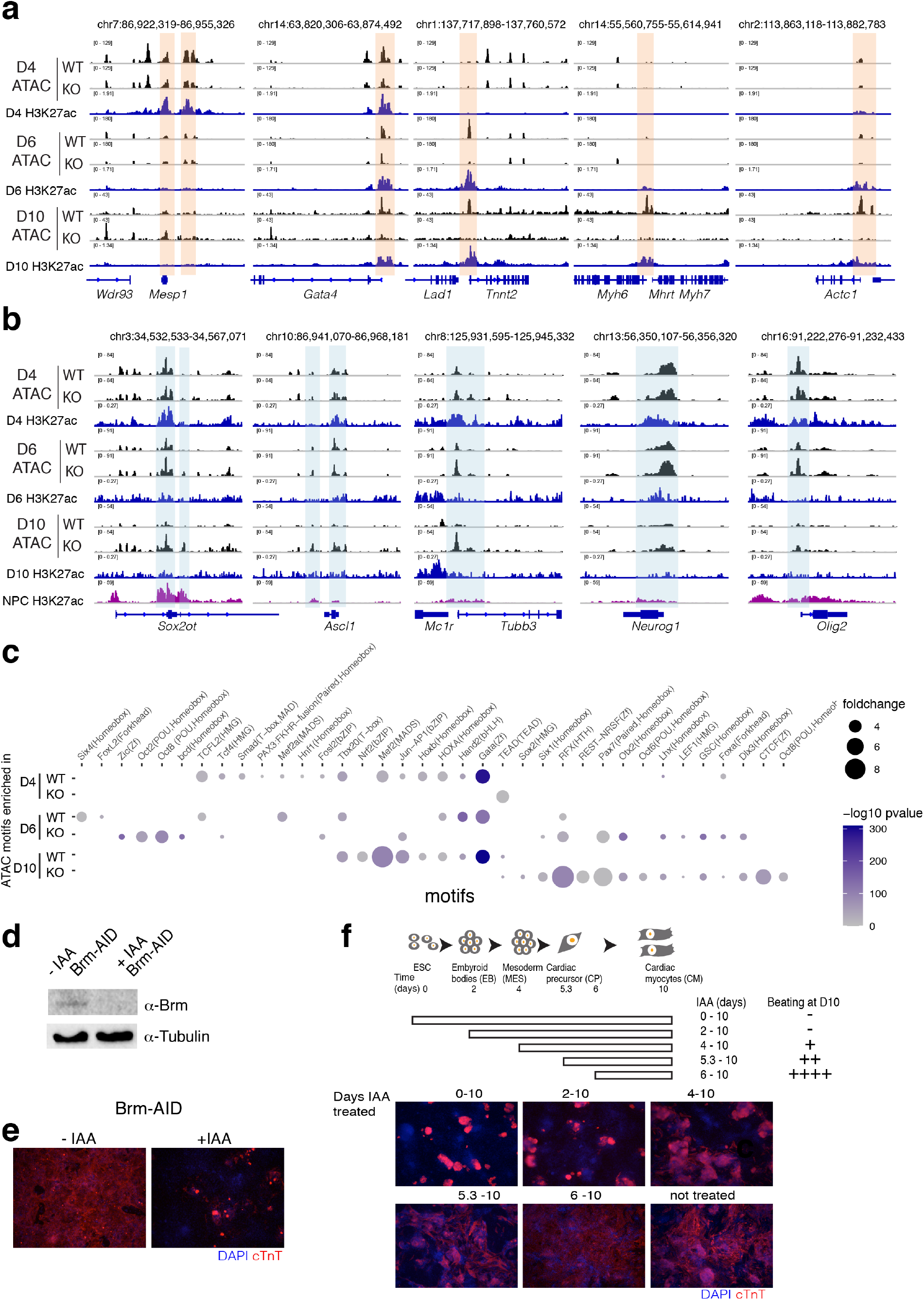
BRM is required during cardiac mesoderm formation. **a-b**, ATAC-seq browser tracks showing WT and BRM KO chromatin accessibility at D4, D6 and D10 of cardiac differentiation along with H3K27ac active enhancer marks at each of these stages for a set of cardiac gene loci (**a**) and neural gene loci, along with neural precursor H3K27ac marks (**b**). **c**, Motifs enriched at the open chromatin regions in WT and BRM KO cells at D4, D6, D10 differentiation stages. **d-f**, Auxin inducible degron mouse ES strain of BRM (Brm-AID) with or without auxin analog indole acetic acid (IAA) present throughout cardiac differentiation shows BRM degradation by western blot (**d**), loss of cardiac myocyte by cTnT immunostaining (**e**) and determines D0 to D4 as the window of BRM activity (**f**).

**Extended Data Fig. 5.**
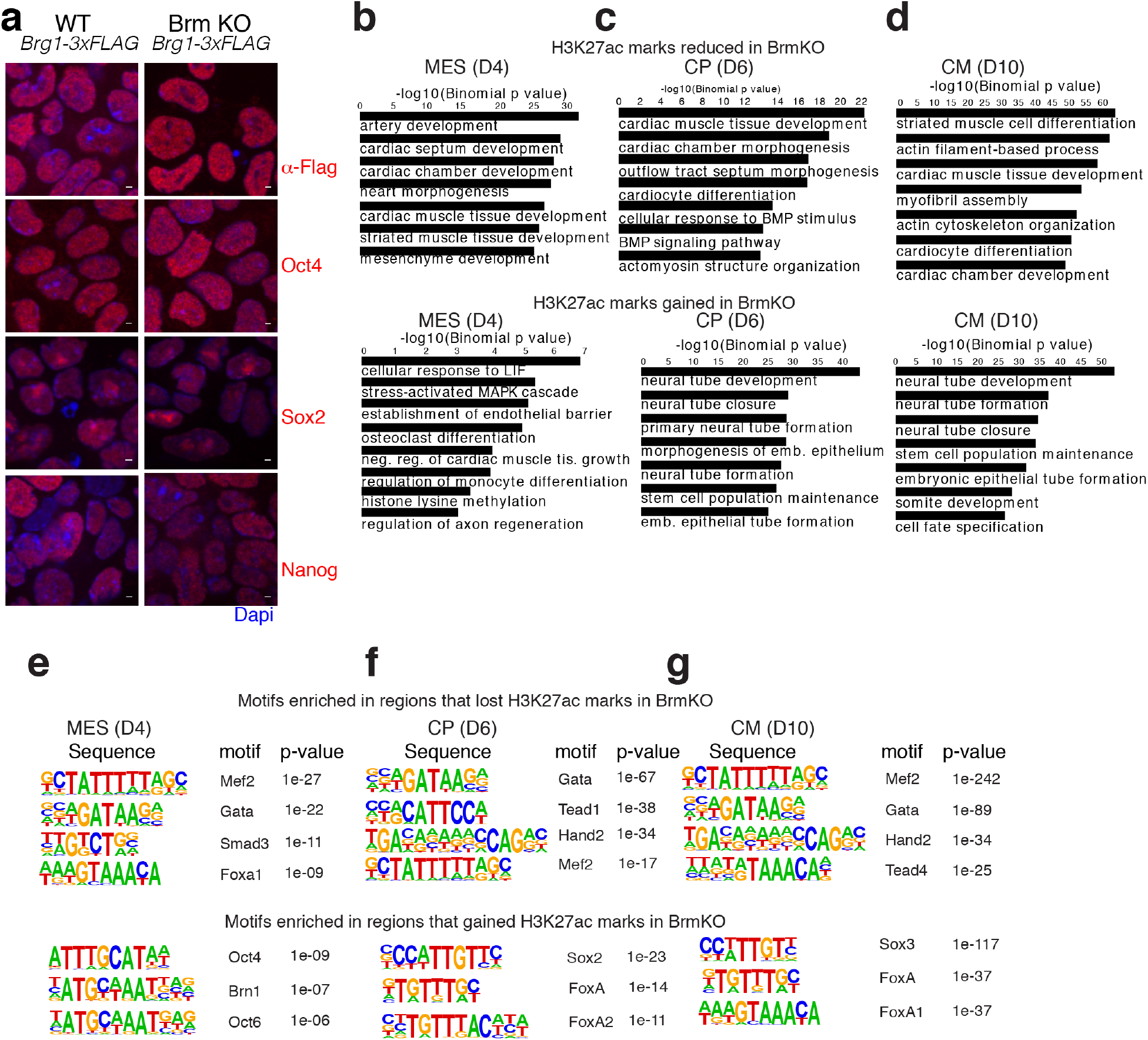
BRM loss leads to reduced H3K27ac marks near cardiac and increased H3K27ac marks near neural genes. **a**, Immunostaining of WT and BRM KO ES cells with indicated pluripotency markers. Scale bars are 2μM, magnification 63x. **b-d**, GO biological processes enriched for genes (within 1mb) near sites that reduced (upper panels) or gained (lower panels) H3K27ac marks in BRM KO cells at D4 (**b**), D6 (**c**) and D10 (**d**) of differentiation. **e-g**, Motifs enriched at the differentially enriched sites in BRM KO cells are shown at D4 (**e**), D6 (**f**) and D10 (**g**) stages of cardiac differentiation respectively.

**Extended Data Fig. 6.**
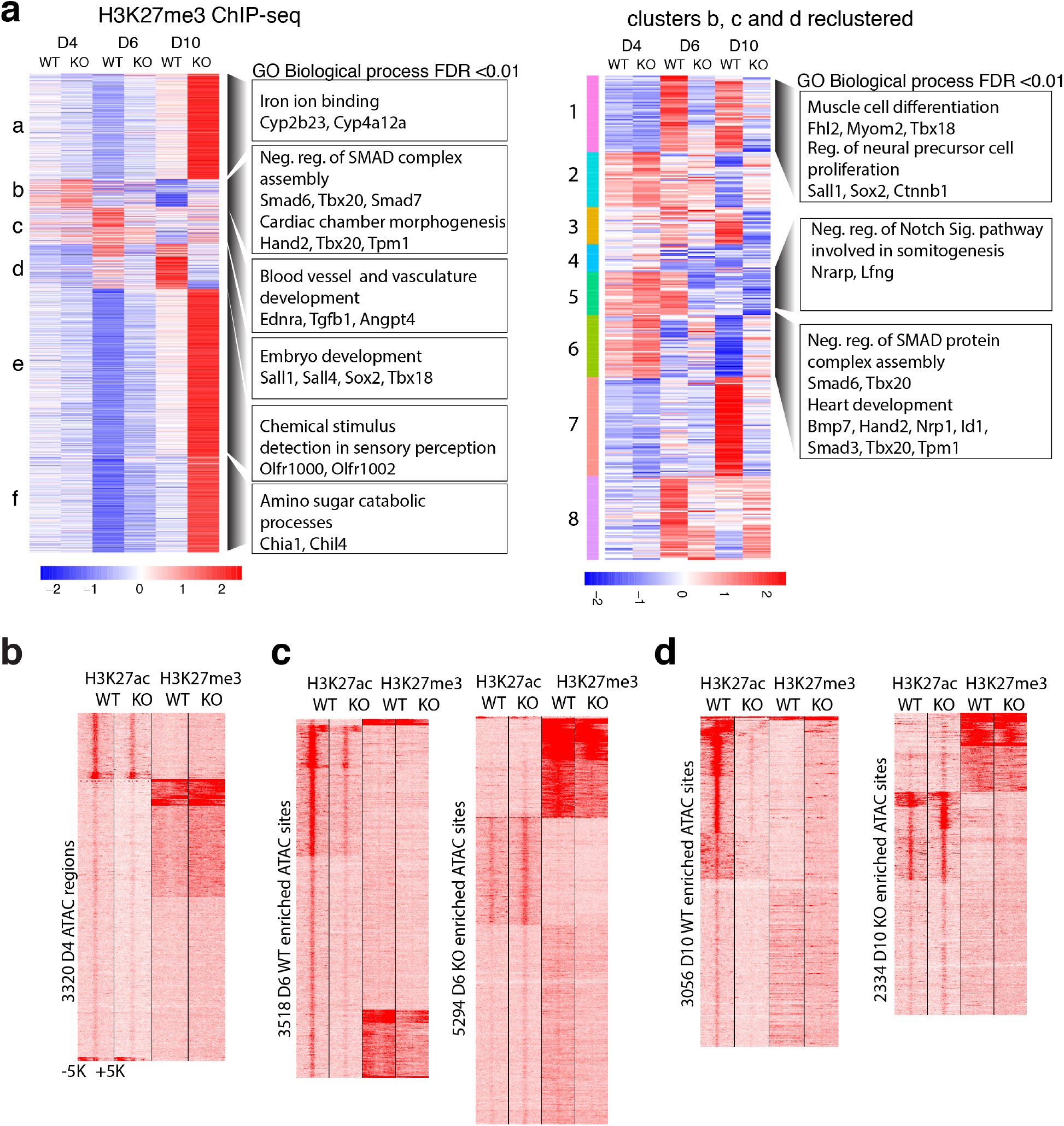
H3K27me3 marks are altered in BRM KO cells at D6 and D10 of differentiation. **a**, Differential enrichment of H3K27me3 marks in WT and BRM KO cells during cardiac differentiation displayed in the form of a heat map. Cluster b, c, and d were re-clustered and shown in a separate heat map (right) GREAT analysis of significant (Benjamini-Hochberg adjusted p-value (FDR) <0.01) GO biological processes (within 1Mb) enrichment for the clusters are on the right with representative genes shown. **b-d**, Correlation of ATAC-seq peaks to active enhancer (H3K27ac) and Polycomb mediated repression (H3K27me3) at D4 (**b**), D6 (**c**) and D10 (**d**) of differentiation.

**Extended Data Fig. 7.**
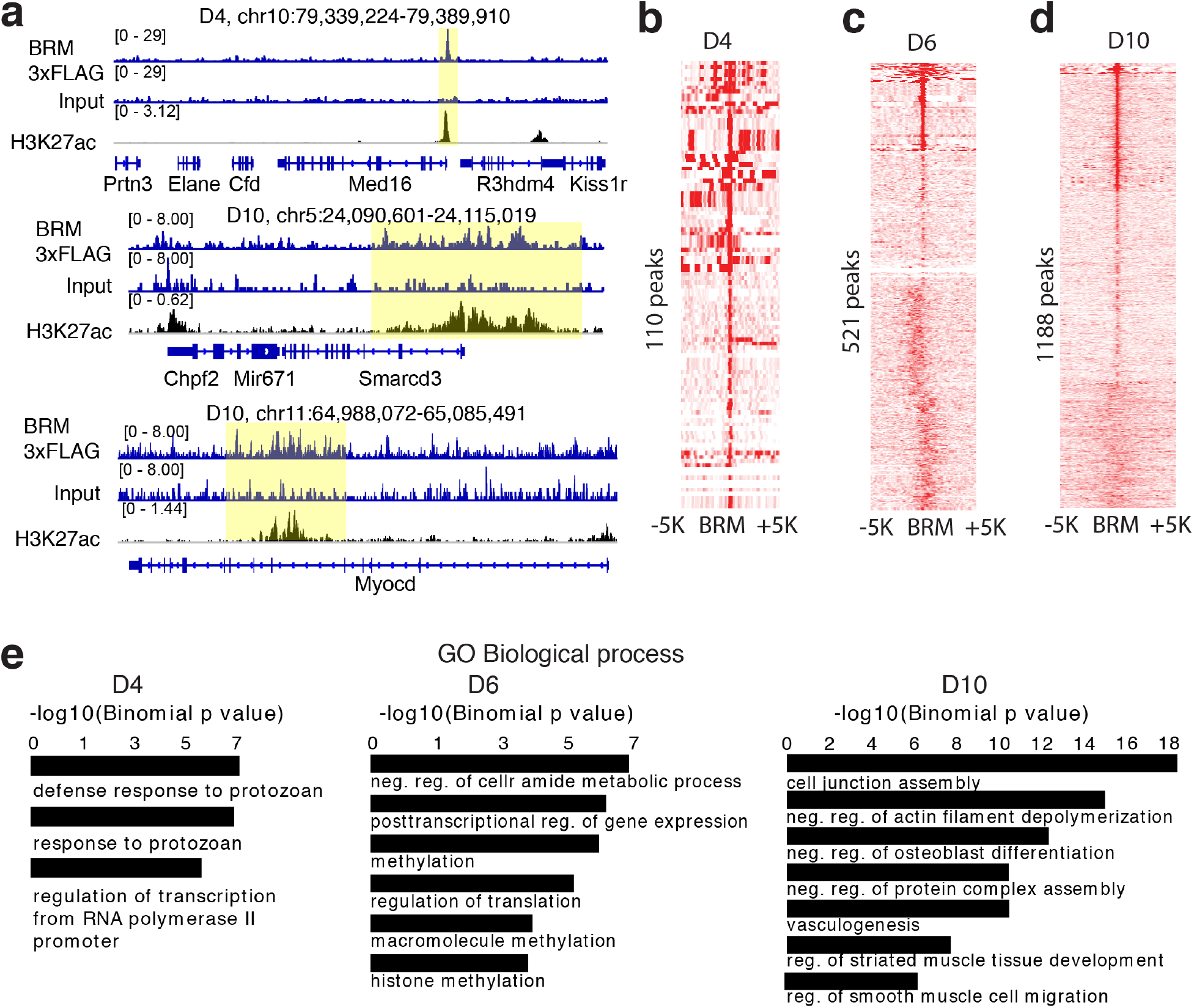
ChIP-seq reveals increased BRM binding during cardiac differentiation. **a-d**, Genome browser (IGV) tracks showing BRM-3xFLAG ChIP-seq, corresponding input samples and activating enhancer mark H3K27ac (**a**) and heat maps of BRM-3xFLAG ChIP-seq over identified BRM binding sites at D4 (**b**), D6 (**c**) and D10 (**d**) of differentiation. **e**, GO biological processes enriched for BRM binding site (within 100kb) at the indicated differentiation stages.

**Extended Data Fig. 8.**
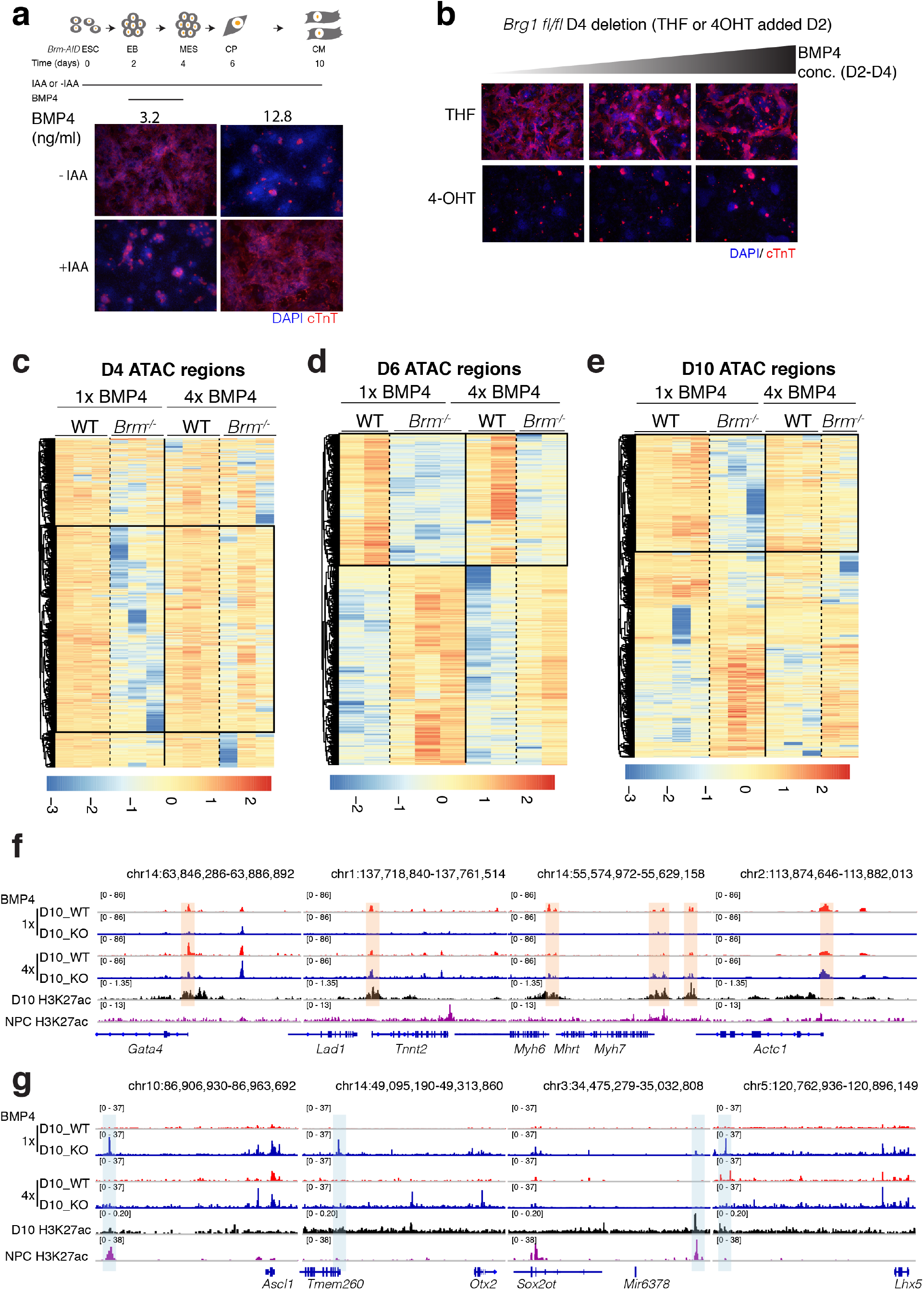
BMP4 restores WT-like chromatin in *Brm*^*−/−*^ cells. **a**, Scheme of cardiac differentiation showing timing of IAA and BMP4 addition. Cardiac troponin T (cTnT) immunostaining of an auxin inducible degron strain of BRM (*Brm-AID*) at D10 of differentiation induced with two different BMP4 concentration with or without IAA. **b**, Immunostaining with cTnT shows that Brg1 loss is not rescued by addition of increasing amount of BMP4. **c-e**, Heat maps showing differential enrichment of ATAC-seq peaks of WT and BRM KO cells at D4 (**c**), D6(**d**) and D10 (**e**) of cardiac differentiation with normal (1x) and high (4x) BMP4 concentrations. Boxed regions show restoration of WT-like chromatin in KO cells at high BMP4 condition. Vertical lanes show replicate data. **f-g**, Browser tracks show chromatin accessibility in WT and BRM KO cells along with H3K27ac marks in cardiomyocytes and neural precursor cells (purple track) near indicated cardiac genes (**f**) and neural genes (**g**)

**Extended Data Fig. 9.**
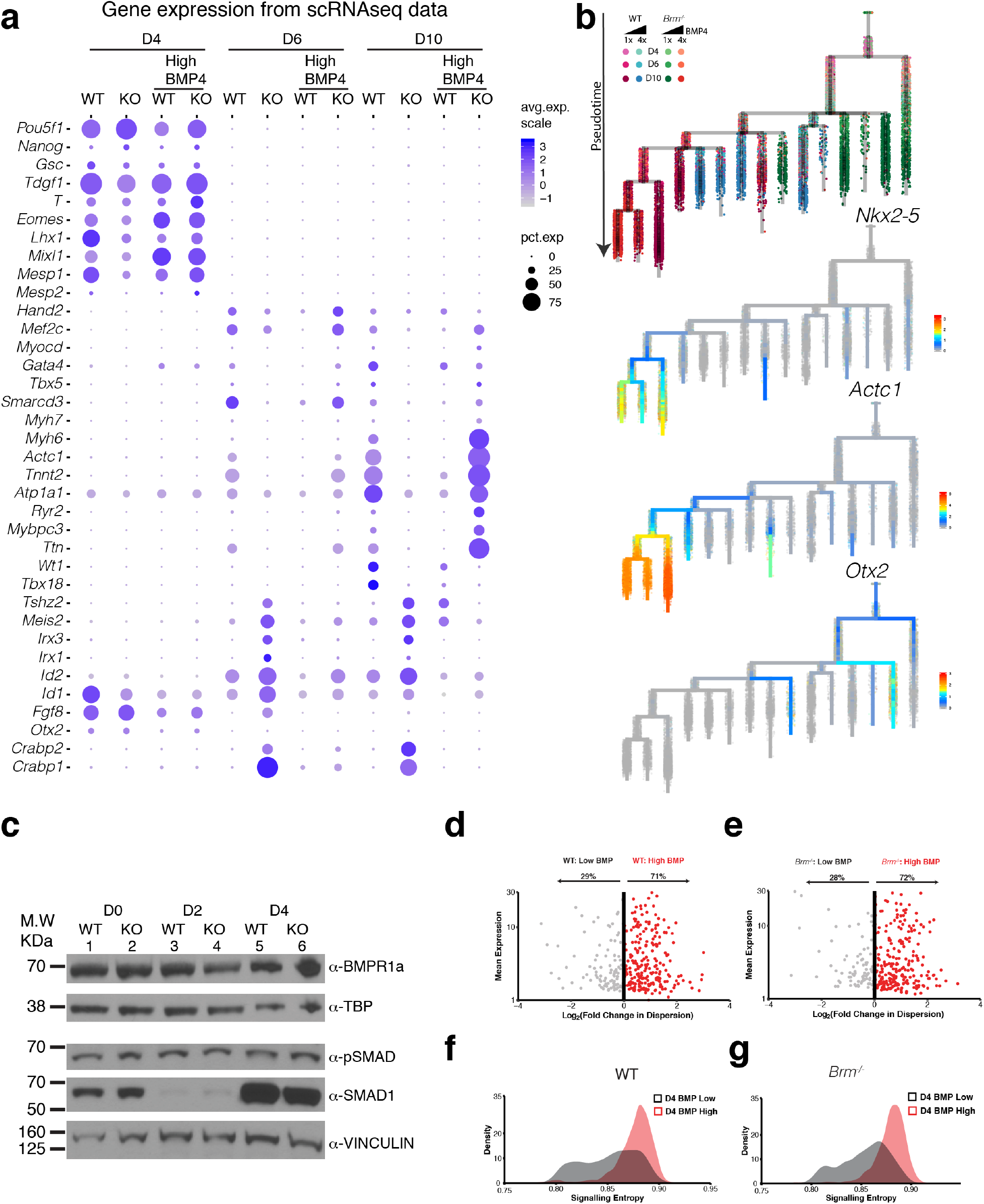
BMP4 increases gene expression noise to restore WT-like gene expression in *Brm*^*−/−*^ cells. **a**, Dot plots showing quantitative changes in gene expression between WT and BRM KO cells induced with normal (1x) or high (4x) BMP4 concentrations at D4, D6 and D10 stages of differentiation for early developmental, cardiac mesoderm, precursors, myocytes and genes enriched in BRM KO cells. **b**, Transcriptional trajectory analysis of WT and BRM KO cells in presence of normal or high BMP4 concentrations showing the genotype representation (top) and URD feature plots of expression of *Nkx2-5*, *Actc1* and *Otx2.* **c**, Western blots showing BMP receptor, Smad1 and phosphor-SMAD expression during D0 to D4 of cardiac differentiation, **d-e**, Scatter plots of single cell RNASeq data showing mean gene expression and variance from mean gene expression at D4 stage of differentiation for WT (**d**) and BRM KO cells (**e**) in low and high BMP4 conditions. **f-g**, Signaling entropy calculated similarly for WT (**f**) and BRM KO cells (**g**) with low and high BMP4 conditions.

**Extended Data Fig. 10.**
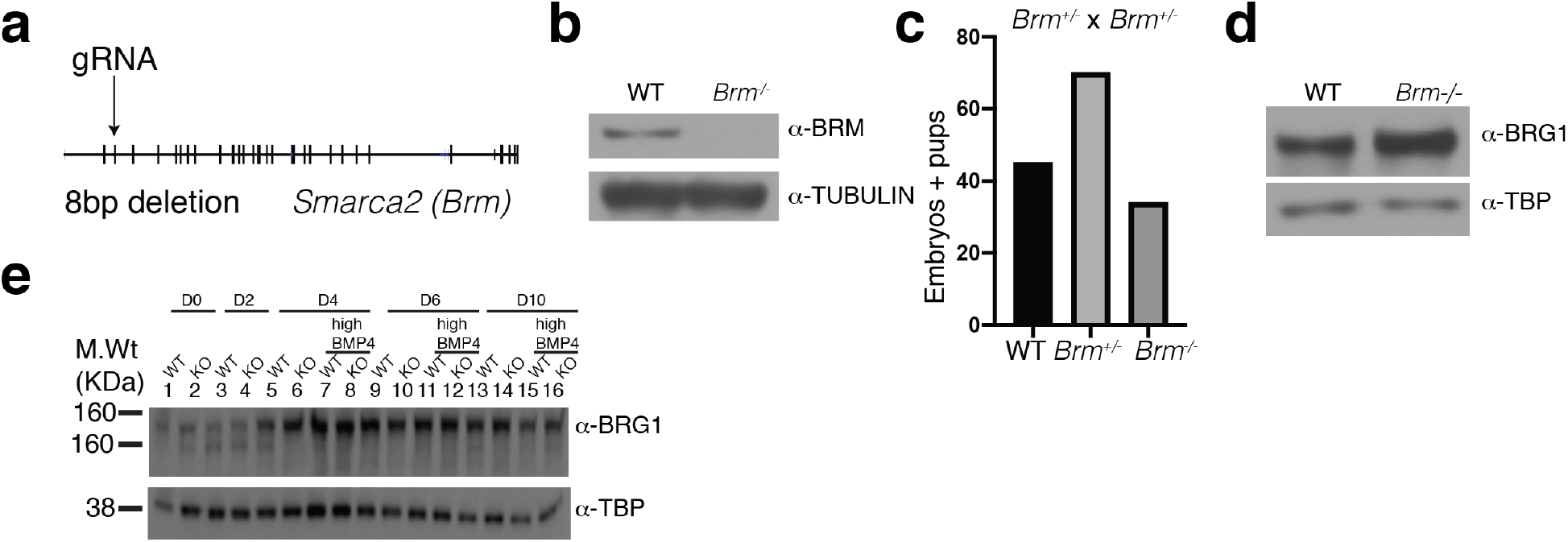
BRG1 partially compensates for BRM loss *in-vivo*. **a**, The exon–intron organization of *Smarca2* (encodes BRM) and the site of guide RNA that targets exon2. The mouse strain from this transfection had an 8 bp deletion leading to premature stop codon. **b**, Western blot with anti-BRM antibody showing loss of BRM protein in Brm-/-mouse brain whole cell extract. a-tubulin is used as a loading control. **c**, Heterozygous Brm mouse mating resulted in pups and embryos at a mendelian ratio. **d-e**, Western blot with antibody against BRG1 shows partial BRG1 compensation in absence of BRM in mouse brain (d), but no compensation in the in-vitro cardiac differentiation system (e).

## References

1. Waddington, C. H. Canalization of Development and the Inheritance of Acquired Characters. Nature 150, 563–565 (1942).

2. Waddington, C. H. The Strategy of the Genes, a Discussion of Some Aspects of Theoretical Biology, by C. H. Waddington,… With an Appendix [Some Physico-chemical Aspects of Biological Organisation] by H. Kacser,… (1957).

3. Ferrell, J. E., Jr. Bistability, Bifurcations, and Waddington’s Review Epigenetic Landscape. Current Biology 22, R458–R466 (2012).

4. Moris, N., Pina, C. & Arias, A. M. Transition states and cell fate decisions in epigenetic landscapes. Nature Publishing Group 17, 693–703 (2016).

5. Hota, S. K. et al. Dynamic BAF chromatin remodeling complex subunit inclusion promotes temporally distinct gene expression programs in cardiogenesis. Development 146, dev174086 (2019).

6. Reyes, J. C. et al. Altered control of cellular proliferation in the absence of mammalian brahma (SNF2alpha). The EMBO Journal 17, 6979–6991 (1998).

7. Van Houdt, J. K. J. et al. Heterozygous missense mutations in SMARCA2 cause Nicolaides-Baraitser syndrome. Nature Publishing Group 1–6 (2012). doi:10.1038/ng.1105

8. Tsurusaki, Y. et al. Mutations affecting components of the SWI/SNF complex cause Coffin-Siris syndrome. Nat Genet 44, 376–378 (2012).

9. Albini, S. et al. Brahma is required for cell cycle arrest and late muscle gene expression during skeletal myogenesis. EMBO reports 16, 1037–1050 (2015).

10. Wu, J. et al. Inactivation of SMARCA2 by promoter hypermethylation drives lung cancer development. Gene 687, 193–199 (2019).

11. Zhang, Z. et al. BRM/SMARCA2 promotes the proliferation and chemoresistance of pancreatic cancer cells by targeting JAK2/STAT3 signaling. Cancer Letters 402, 213–224 (2017).

12. Willis, M. S. et al. BRG1 and BRM function antagonistically with c-MYC in adult cardiomyocytes to regulate conduction and contractility. J. Mol. Cell. Cardiol. 105, 99–109 (2017).

13. Januario, T. et al. PRC2-mediated repression of SMARCA2 predicts EZH2 inhibitor activity in SWI/SNF mutant tumors. Proc. Natl. Acad. Sci. U.S.A. 114, 12249–12254 (2017).

14. Smith-Roe, S. L. & Bultman, S. J. Combined gene dosage requirement for SWI/SNF catalytic subunits during early mammalian development. Mamm. Genome 24, 21–29 (2013).

15. Hoffman, G. R. et al. Functional epigenetics approach identifies BRM/SMARCA2 as a critical synthetic lethal target in BRG1-deficient cancers. Proc. Natl. Acad. Sci. U.S.A. 111, 3128–3133 (2014).

16. Wiley, M. M., Muthukumar, V., Griffin, T. M. & Griffin, C. T. SWI/SNF chromatin-remodeling enzymes Brahma-related gene 1 (BRG1) and Brahma (BRM) are dispensable in multiple models of postnatal angiogenesis but are required for vascular integrity in infant mice. J Am Heart Assoc 4, e001972–e001972 (2015).

17. McInnes, L., Healy, J. & Melville, J. UMAP: Uniform Manifold Approximation and Projection for Dimension Reduction. arXiv.org stat.ML, arXiv:1802.03426 (2018).

18. Wolf, F. A. et al. PAGA: graph abstraction reconciles clustering with trajectory inference through a topology preserving map of single cells. Genome Biol. 20, 59–9 (2019).

19. Farrell, J. A. et al. Single-cell reconstruction of developmental trajectories during zebrafish embryogenesis. Science 360, eaar3131 (2018).

20. Edri, S., Hayward, P., Jawaid, W., Development, A. A.2019. Neuro-mesodermal progenitors (NMPs): a comparative study between pluripotent stem cells and embryo-derived populations. Development doi:10.1242/dev.180190.supplemental

21. Gouti, M. et al. A Gene Regulatory Network Balances Neural and Mesoderm Specification during Vertebrate Trunk Development. 1–27 (2017). doi:10.1016/j.devcel.2017.04.002

22. Pankratz, M. T. et al. Directed Neural Differentiation of Human Embryonic Stem Cells via an Obligated Primitive Anterior Stage. Stem Cells 25, 1511–1520 (2007).

23. Thomson, M. et al. Pluripotency Factors in Embryonic Stem Cells Regulate Differentiation into Germ Layers. Cell 145, 875–889 (2011).

24. Jang, S. et al. Dynamics of embryonic stem cell differentiation inferred from single-cell transcriptomics show a series of transitions through discrete cell states. Elife 6, 91 (2017).

25. Alexander, J. M. et al. Brg1 modulates enhancer activation in mesoderm lineage commitment. Development 142, 1418–1430 (2015).

26. Takeuchi, J. K. et al. Chromatin remodelling complex dosage modulates transcription factor function in heart development. Nature Communications 2, 187–11 (2011).

27. Ho, L. et al. An embryonic stem cell chromatin remodeling complex, esBAF, is essential for embryonic stem cell self-renewal and pluripotency. Proc. Natl. Acad. Sci. U.S.A. 106, 5181–5186 (2009).

28. Kadam, S. & Emerson, B. M. Transcriptional specificity of human SWI/SNF BRG1 and BRM chromatin remodeling complexes. Molecular Cell 11, 377–389 (2003).

29. Raab, J. R., Runge, J. S., Spear, C. C. & Magnuson, T. Co-regulation of transcription by BRG1 and BRM, two mutually exclusive SWI/SNF ATPase subunits. Epigenetics Chromatin 1–15 (2017). doi:10.1186/s13072-017-0167-8

30. Corces, M. R. et al. An improved ATAC-seq protocol reduces background and enables interrogation of frozen tissues. Nat Meth 14, 959–962 (2017).

31. Wamstad, J. A. et al. Dynamic and coordinated epigenetic regulation of developmental transitions in the cardiac lineage. Cell 151, 206–220 (2012).

32. Rhee, H. S. et al. Expression of Terminal Effector Genes in Mammalian Neurons Is Maintained by a Dynamic Relay of Transient Enhancers. Neuron 92, 1252–1265 (2016).

33. Nora, E. P. et al. Targeted Degradation of CTCF Decouples Local Insulation of Chromosome Domains from Genomic Compartmentalization. Cell 169, 930–944.e22 (2017).

34. Alver, B. H. et al. The SWI/SNF chromatin remodelling complex is required for maintenance of lineage specific enhancers. Nature Communications 8, 14648 (2017).

35. Li, C. et al. Concerted genomic targeting of H3K27 demethylase REF6 and chromatin-remodeling ATPase BRM in Arabidopsis. Nat Genet 48, 687–693 (2016).

36. Tie, F., Banerjee, R., Conrad, P. A., Scacheri, P. C. & Harte, P. J. Histone Demethylase UTX and Chromatin Remodeler BRM Bind Directly to CBP and Modulate Acetylation of Histone H3 Lysine 27. Molecular and Cellular Biology 32, 2323–2334 (2012).

37. Zhu, Q. et al. The transcription factor Pou3f1 promotes neural fate commitment via activation of neural lineage genes and inhibition of external signaling pathways. Elife 3, 2–21 (2014).

38. Kattman, S. J. et al. Stage-specific optimization of activin/nodal and BMP signaling promotes cardiac differentiation of mouse and human pluripotent stem cell lines. Cell Stem Cell 8, 228–240 (2011).

39. Paulsen, M., Legewie, S., Eils, R., Karaulanov, E. & Niehrs, C. Negative feedback in the bone morphogenetic protein 4 (BMP4) synexpression group governs its dynamic signaling range and canalizes development. in 108, 10202–10207 (National Academy of Sciences, 2011).

40. Arias, A. M. & Hayward, P. Filtering transcriptional noise during development: concepts and mechanisms. Nat Rev Genet 7, 34–44 (2006).

41. Bier, E. & De Robertis, E. M. EMBRYO DEVELOPMENT. BMP gradients: A paradigm for morphogen-mediated developmental patterning. Science 348, aaa5838–aaa5838 (2015).

42. Eling, N., Richard, A. C., Richardson, S., Marioni, J. C. & Vallejos, C. A. Correcting the Mean-Variance Dependency for Differential Variability Testing Using Single-Cell RNA Sequencing Data. Cell Systems 7, 284–294.e12 (2018).

43. Enver, T. & Teschendorff, A. E. Single-cell entropy for accurate estimation of differentiation potency from a cell’s transcriptome. Nature Communications 8, 1–15 (2017).

44. Thompson, K. W., Marquez, S. B., Lu, L. & Reisman, D. Induction of functional Brm protein from Brm knockout mice. Oncoscience 2, 349–361 (2015).

45. Dupin, E., Calloni, G. W., Coelho-Aguiar, J. M. & Le Douarin, N. M. The issue of the multipotency of the neural crest cells. Developmental Biology 444 Suppl 1, S47–S59 (2018).

46. Motohashi, T. & Kunisada, T. Extended multipotency of neural crest cells and neural crest-derived cells. Curr. Top. Dev. Biol. 111, 69–95 (2015).

47. Srivastava, D. & DeWitt, N. In Vivo Cellular Reprogramming: The Next Generation. Cell 166, 1386–1396 (2016).

48. Tursun, B., Patel, T., Kratsios, P. & Hobert, O. Direct conversion of C. elegans germ cells into specific neuron types. Science 331, 304–308 (2011).

49. Cheloufi, S. et al. The histone chaperone CAF-1 safeguards somatic cell identity. Nature 528, 218–224 (2015).

50. Kolundzic, E. et al. FACT Sets a Barrier for Cell Fate Reprogramming in Caenorhabditis elegans and Human Cells. Developmental Cell 46, 611–626.e12 (2018).

51. Jiang, Z. et al. Knockdown of Brm and Baf170, Components of Chromatin Remodeling Complex, Facilitates Reprogramming of Somatic Cells. Stem Cells and Development 24, 2328–2336 (2015).

52. Lalit, P. A. et al. Lineage Reprogramming of Fibroblasts into Proliferative Induced Cardiac Progenitor Cells by Defined Factors. Cell Stem Cell 18, 354–367 (2016).

53. Singhal, N., Esch, D., Stehling, M. & Schöler, H. R. BRG1 Is Required to Maintain Pluripotency of Murine Embryonic Stem Cells. Biores Open Access 3, 1–8 (2014).

54. Takeuchi, J. K. & Bruneau, B. G. Directed transdifferentiation of mouse mesoderm to heart tissue by defined factors. Nature 459, 708–711 (2009).

55. Treutlein, B. et al. Dissecting direct reprogramming from fibroblast to neuron using single-cell RNA-seq. Nature 534, 391–395 (2016)

## References

56. Ho, L. et al. An embryonic stem cell chromatin remodeling complex, esBAF, is an essential component of the core pluripotency transcriptional network. Proc. Natl. Acad. Sci. U.S.A. 106, 5187–5191 (2009).

57. Conti, L. et al. Niche-Independent Symmetrical Self-Renewal of a Mammalian Tissue Stem Cell. PLoS Biol 3, e283–13 (2005).

58. Cong, L. et al. Multiplex genome engineering using CRISPR/Cas systems. Science 339, 819–823 (2013).

59. Abmayr, S. M., Yao, T., Parmely, T. & Workman, J. L. Preparation of nuclear and cytoplasmic extracts from mammalian cells. Curr Protoc Pharmacol Chapter 12, Unit12.3-12.3.10 (2006).

60. Kim, D. et al. TopHat2: accurate alignment of transcriptomes in the presence of insertions, deletions and gene fusions. Genome Biol. 14, R36 (2013).

61. Liao, Y., Smyth, G. K. & Shi, W. featureCounts: an efficient general purpose program for assigning sequence reads to genomic features. Bioinformatics 30, 923–930 (2014).

62. Robinson, M. D., McCarthy, D. J. & Smyth, G. K. edgeR: a Bioconductor package for differential expression analysis of digital gene expression data. Bioinformatics 26, 139–140 (2010).

63. Zambon, A. C. et al. GO-Elite: a flexible solution for pathway and ontology over-representation. Bioinformatics 28, 2209–2210 (2012).

64. Satija, R., Farrell, J. A., Gennert, D., Schier, A. F. & Regev, A. Spatial reconstruction of single-cell gene expression data. Nat. Biotechnol. 33, 495–502 (2015).

65. Lambiotte, R., Delvenne, J. C. & Barahona, M. Laplacian Dynamics and Multiscale Modular Structure in Networks. arXiv.org physics.soc-ph, arXiv:0812.1770–90 (2008).

66. Langmead, B. & Salzberg, S. L. Fast gapped-read alignment with Bowtie 2. Nat Meth 9, 357–359 (2012).

67. Thomas, R., Thomas, S., Holloway, A. K. & Pollard, K. S. Features that define the best ChIP-seq peak calling algorithms. Brief. Bioinformatics 18, 441–450 (2017).

68. Afgan, E. et al. The Galaxy platform for accessible, reproducible and collaborative biomedical analyses: 2018 update. Nucleic Acids Research 46, W537–W544 (2018).

69. McLean, C. Y. et al. GREAT improves functional interpretation of cis-regulatory regions. Nat. Biotechnol. 28, 495–501 (2010).

70. O’Geen, H., Echipare, L. & Farnham, P. J. Using ChIP-seq technology to generate high-resolution profiles of histone modifications. Methods Mol. Biol. 791, 265–286 (2011).

71. Heinz, S. et al. Simple combinations of lineage-determining transcription factors prime cis-regulatory elements required for macrophage and B cell identities. Molecular Cell 38, 576–589 (2010).

72. Xing, H., Mo, Y., Liao, W. & Zhang, M. Q. Genome-Wide Localization of Protein-DNA Binding and Histone Modification by a Bayesian Change-Point Method with ChIP-seq Data. PLoS Comput Biol 8, e1002613–12 (2012).

73. McCarthy, D. J., Chen, Y. & Smyth, G. K. Differential expression analysis of multifactor RNA-Seq experiments with respect to biological variation. Nucleic Acids Research 40, 4288–4297 (2012).

